# Stochastic Assembly and Metabolic Network Reorganization Drive Microbial Resilience in Arid Soils

**DOI:** 10.1101/2025.02.21.639521

**Authors:** Christian Ayala-Ortiz, Viviana Freire-Zapata, Malak M. Tfaily

## Abstract

Microbial resilience plays a pivotal role in ecosystems as environmental fluctuations impact community functioning and stability. Despite resilience emerging from both individual adaptations and community-level processes, integration of these mechanisms remains enigmatic, particularly in arid environments. These extreme ecosystems, spanning over 45% of Earth’s terrestrial surface, provide a natural laboratory for understanding microbial survival under harsh conditions. Here, we use time-resolved multi-omics to show that resilience results from dynamic microbial network reorganization enabling the coordination between stochastic processes that maintain community stability, and individual stress responses. Additionally, Thermoproteota emerged as a keystone taxon maintaining nitrogen cycling and fostering cross-feeding networks. Its ecological prominence highlights its central role in arid ecosystems, making it an ideal model organism for understanding microbial adaptation to environmental extremes. Our findings bridge the gap between individual adaptations and community-wide resilience, offering a framework for understanding microbial responses to environmental fluctuations and their implications for ecosystem function.

## INTRODUCTION

Microbial communities are pivotal to ecosystem functions ^1^, yet they face persistent environmental fluctuations across diverse habitats ^2^. These changes can dramatically impact community functioning and stability ^3^ with responses typically manifesting through four primary scenarios: full recovery, physiological adaptation, functional redundancy, or complete loss of function^4^. Despite advances, the molecular mechanisms underlying microbial community recovery remain poorly understood, particularly how individual strategies integrate with community-level processes to ensure resilience ^5^.

Microbial resilience emerges from both individual survival strategies and collective community-level coordination ^6–8^. Community-level dynamics like functional redundancy can buffer ecosystem functions against environmental stress ^4^, while individual adaptations provide the mechanistic basis for survival during extreme fluctuations^9–12^. Understanding this multi-level response is particularly crucial as climate change intensifies the frequency and severity of environmental perturbations.

Arid environments offer a unique laboratory for studying microbial resilience.

Characterized by extreme temperatures, intense UV radiation, and variable precipitation ^13,14^, these ecosystems represent critical systems for understanding adaptation ^15,16^. As these regions expand due to climate change ^17,18^, understanding their ecological mechanisms becomes increasingly urgent for both ecosystem stability and agricultural resilience.

Soil microbes have evolved diverse survival strategies to cope with extreme conditions. Dormancy allows bacteria to persist during unfavorable conditions and rapidly reactivate when conditions improve ^19,20^, ensuring long-term population survival ^21^. Upon rewetting, dormant microbes prioritize DNA repair and energy generation ^20^. Other adaptations, such as the consistent transcription of stress response and nutrient acquisition genes during dry periods, further enhance survival ^22^, while mechanisms to mitigate oxidative and osmotic stress protect cellular integrity ^23^. However, how these individual adaptations integrate with community-level processes remains largely unknown, leaving crucial gaps in our understanding of how they scale to maintain ecosystem function, particularly during dramatic environmental transitions like wet-dry cycles ^13^.

To address this knowledge gap, we employed an integrated time-resolved multiomics approach analyzing bare soil samples from Arizona’s Saguaro National Park, an iconic desert ecosystem characterized by extreme wet-dry cycles. Our findings reveal that microbial resilience emerges from a sophisticated balance between community stability and functional adaptability. While stochastic processes maintain stable microbial community composition, the organic matter profiles show deterministic shifts reflecting metabolic adaptations to environmental stress. The key mechanism enabling this dual stability-adaptability strategy is microbial network reorganization, allowing communities to maintain both compositional integrity and coordinated individual responses. This dynamic is exemplified by ammonia-oxidizing Thermoproteota, whose flexible gene expression patterns enable it to assume different network roles while preserving critical ecosystem functions during environmental fluctuations. Our findings advance the understanding of microbial resilience in expanding drylands and provide a framework for predicting ecosystem responses to climate change. By comprehensively characterizing both microbial communities and their metabolic outputs, we demonstrate how coordinated responses across multiple organizational levels maintain biogeochemical cycling under extreme conditions—insights critical for ecosystem management as arid regions continue to expand globally.

## RESULTS

### Monsoon events create distinct physicochemical states in arid soils

The North American Monsoon drives dramatic shifts in resource availability across desert ecosystems. We characterized these environmental transitions through intensive temporal sampling at the Saguaro National Park (Fig. 1a). Soil samples were collected at 10 cm depth across eight time points from May to September 2021, with an additional sampling in May 2022 (n = 36, 4 samples per time point), capturing the complete monsoon cycle from pre-monsoon drought through post-monsoon recovery (Fig. 1b). While these environmental transitions aligned with historical monsoon patterns in the Southwestern USA ^26^, 2021 saw some of the highest precipitation levels of the last decade (Supplementary Fig. 1a), offering an ideal natural experiment to examine microbial responses to extreme environmental transitions.

**Figure 1.**
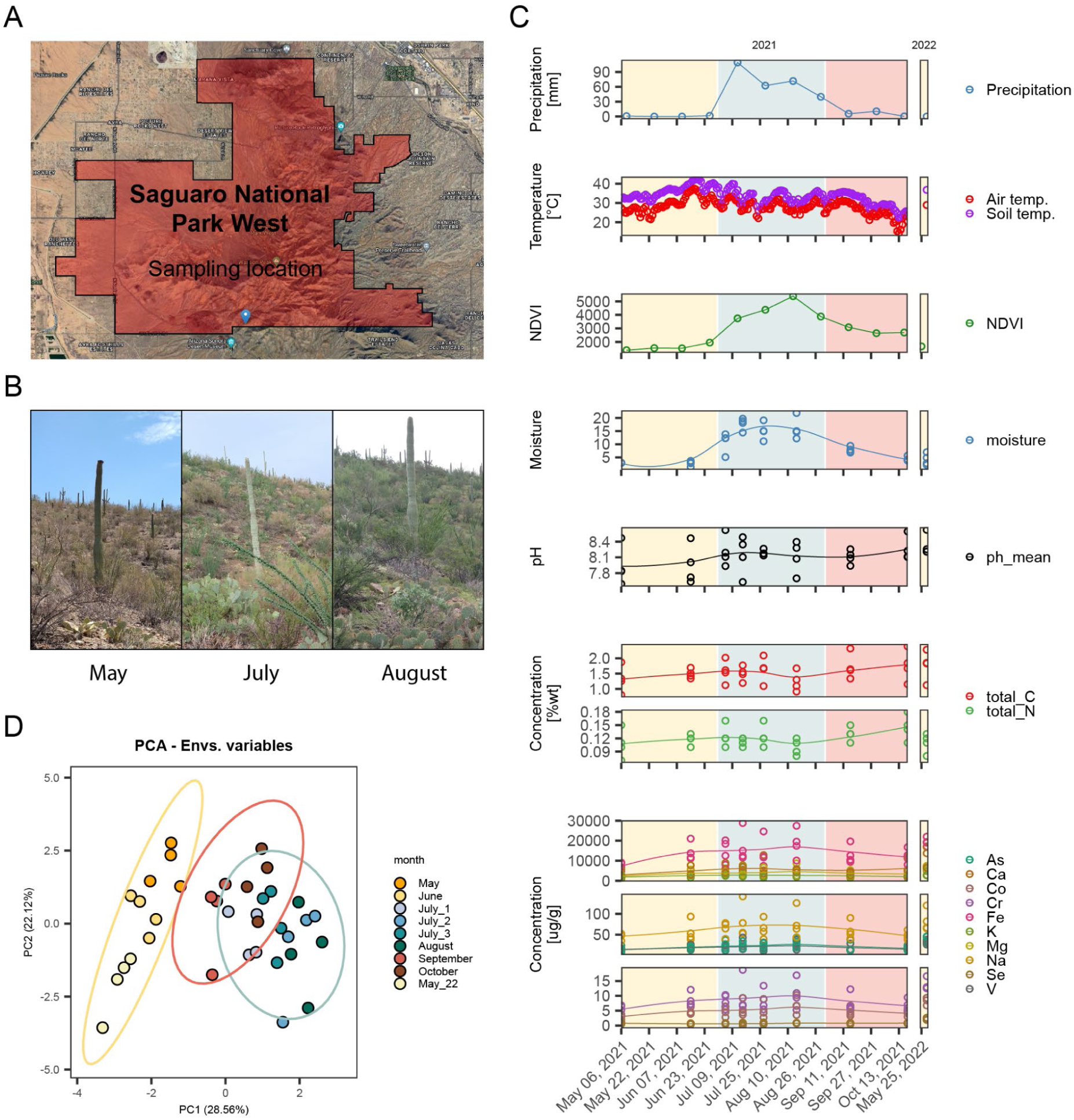
(A) Geographic location of soil sampling sites within Saguaro National Park. (B) Pictures taken at the sampling site during the sampling campaign in May, July and August of 2021. (C) Principal ordination analysis of the collected samples (n = 36) based on the measured environmental factors. (D) Temporal dynamics of environmental and physicochemical factors throughout the 2021 monsoon season.

The shift from extended drought to intense rainfall events created distinct temporal patterns in soil physicochemical conditions, marked by substantial changes in moisture availability, temperature, and potential resource inputs (Fig. 1c, Supplementary Table 1, Supplementary Fig. 1b). Principal component analysis revealed three distinct sample clusters corresponding to pre-monsoon (yellow ellipse), monsoon (blue ellipse), and post-monsoon (red ellipse) periods (Fig. 1d). Notably, post-monsoon samples gradually returned to conditions similar to the pre-monsoon state, highlighting the cyclical nature of these environmental transitions. Moisture content and vegetation density (measured by the Normalized Difference Vegetation Index - NDVI) emerged as primary drivers, explaining 30.47% of observed variance along PC1, while air temperature accounted for 19.66% along PC2. Hierarchical clustering analysis of physicochemical parameters further confirmed these distinct environmental states between monsoon and non-monsoon periods (Supplementary Fig. 1b).

### Community-level responses emerge from local adaptation

To understand microbial responses to monsoon transitions, we employed a multi-omics approach that combined microbial taxonomic profiling (using ASVs derived from 16S rRNA amplicon sequencing and OTUs derived from shotgun metagenomics) with organic matter characterization using Fourier-transform ion cyclotron resonance mass spectrometry, (FTICR-MS).

Our analysis revealed microbial compositional stability despite environmental fluctuations (Fig. 2a, c-d, Supplementary Fig. 2a; Supplementary Note 1). Beta diversity analysis showed only slight community differences during monsoon months (PERMANOVA, p = 0.033), with communities returning to their original structure after the storms (Fig 2d, Extended Fig 2b). This resilience aligns with prior studies suggesting arid soil microbial communities are adapted to endure dry-wet cycles ^27^ through ecological memory ^28^, optimized for historical moisture regimes ^29^. However, alpha diversity metrics presented a more dynamic response (Fig. 2f). Unlike previous reports of stable or increased microbial diversity post-precipitation ^16,30^, we observed a temporary decline in microbial diversity followed by recovery. This decline likely results from mechanisms such as differential growth triggered by increased water availability ^31^, formation of anoxic microsites ^32^, and osmotic stress-induced mortality ^33^. Comparison of metagenomics read signatures with other global arid ecosystems revealed that while each ecosystem harbors locally adapted communities, Sonoran Desert more closely resembled those from North American deserts (e.g., Mojave) than geographically distant systems (e.g., Negev) (Fig.2c) suggesting local adaptation shapes community composition while shared regional species pools support resilience ^34,35^.

**Figure 2.**
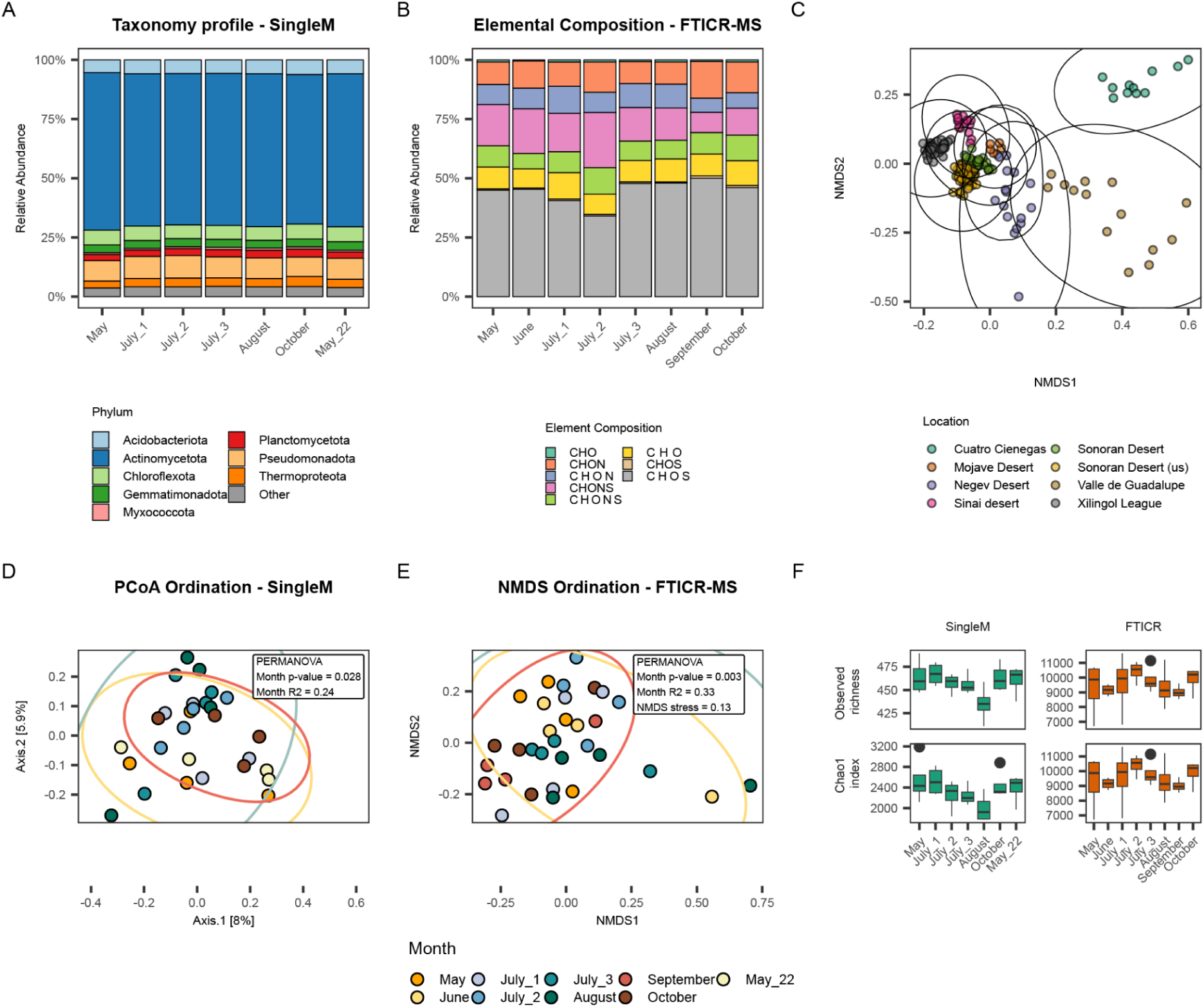
Microbial community and organic matter profiling of arid soil samples. (A) Average relative abundance of bacterial phyla, from OTUs inferred with SingleM, at each sampling time (n = 4). (B) Average relative abundance of observed elemental compositions based on FTICR-MS data collected in positive mode at each sampling time (n = 4). (C) Ordination of different metagenomics datasets collected from arid soils. (D) Principal component analysis (PCoA) of shotgun metagenomics data (n = 28). (E) Non-metric multidimensional scaling (NDMS) of FTICR-MS data (n = 28). (F) Alpha diversity indices (observed richness and Chao1 richness) inferred from SingleM OTUs and FTICR-MS metabolites abundances respectively. Boxes represent the upper and lower quartiles, the line in each box represents the median and the whiskers represent the maximum and minimum values, no further than 1.5 times the interquartile range; values beyond the whiskers represent outliers and are plotted individually

FTICR-MS analysis detected 20,971 different masses (m/z) across all samples, with 4,175 assigned putative molecular formulas. Unlike lignin-dominated peatlands ^36,37^, these arid soils showed predominance of lipid-like compounds, followed by protein- and lignin-like metabolites (Supplementary Fig. 2d; Supplementary Note 1), indicating strong microbial influence on soil organic matter composition ^38^.

Organic matter composition showed significant temporal variation (p = 0.014) (Fig. 2b, e), particularly in N- and S-containing compounds (e.g., CHON, CHONS, CHONP) (Supplementary Fig. 2f, h). During wet periods, osmotic stress-induced cell lysis released intracellular metabolites ^33^, while increased substrate accessibility enhanced microbial activity ^39^. Though some lipid-like compounds remained stable throughout the monsoon season, amino sugar abundance increased with water availability. These amino sugars, likely derived from microbial components ^40^ may have been accumulated due to osmotic stress-induced mortality^32^. The subsequent increase in carbohydrates by the end of the monsoon season (September and August) likely reflected post-monsoon vegetation growth and enhanced microbial activity ^41^. Organic matter alpha diversity initially increased at monsoon onset, likely due to precipitation-induced release of labile metabolites ^42^. These metabolites were subsequently consumed, leading to decreased diversity and liability (NOSC) by monsoon’s end (Fig. 2f, Supplementary Fig. 2e).

### Contrasting assembly processes reveal mechanisms of community-level resilience

While our observations demonstrate clear community-level responses to environmental change, the underlying ecological processes maintaining stability remain unclear. To investigate these mechanisms, we analyzed the ecological processes driving both community assembly and metabolite dynamics ^43–46^.

Our analysis revealed contrasting yet complementary mechanisms: stochastic processes predominantly shaped microbiome assemblages, while deterministic processes governed metabolite assemblages (Fig. 3 a, b). However, the relative importance of these ecological processes shifted dynamically with changing environmental conditions (Fig. 3c).

**Figure 3.**
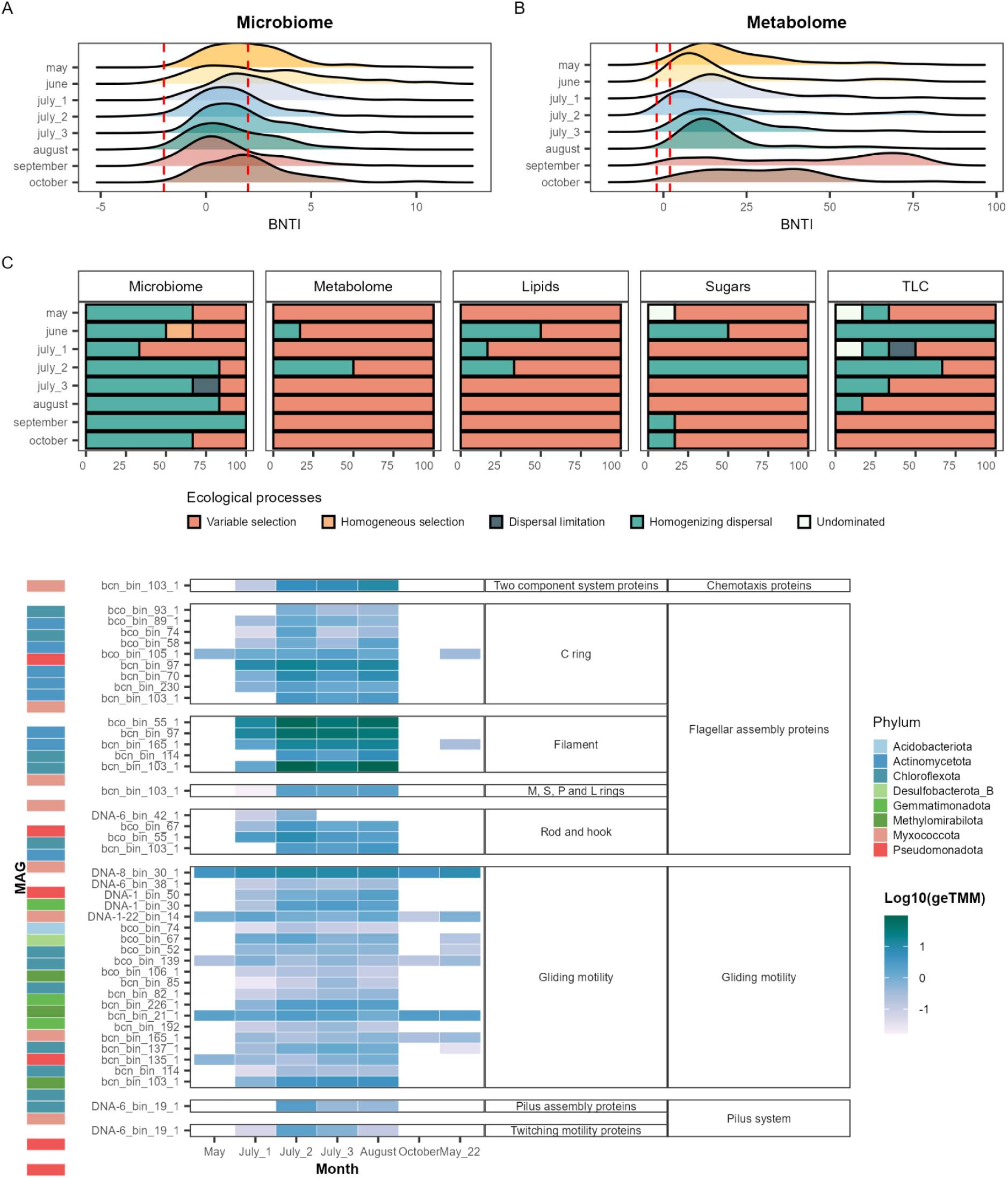
(A) and (B) Density plots showing changes in the βNTI of microbial community and metabolome in each month of sampling (n total = 32), βNTI was calculated from 16s rRNA OTUs and TWCD metabolites. Assembly processes can be delimited by the red dashed lines: variable selection (βNTI > 2), homogenous selection (βNTI < −2), stochastic assembly (|βNTI| < 2). (C) Bar plots showing the putative influence of different ecological processes within each month according to the microbial community, bulk metabolites, lipid-, sugar- and TLC-like metabolites. (D), Heatmap representing the expression of microbial motility related genes. Gene length corrected trimmed mean of M- values (geTMM) values for expressed motility related genes were averaged per each MAG per month, the log10 is represented.

Early monsoon onset (early July) triggered a brief period of deterministic microbial community assembly through environmental filtering and biotic processes. The sudden influx of water and nutrients created selection pressures favoring wet-adapted taxa ^47^ and benefiting tolerant or opportunistic groups ^8^, leading to competition between rain-activated and dry-tolerant microbes ^32^. As rainfall events increased in frequency (late July-early August, Fig. 1c) stochastic processes emerged as the dominant driver of community assembly, maintaining compositional stability (Fig. 3c). This stability emerged from two key mechanisms: enhanced pore connectivity enabled active microbial movement ^4^, while microbial dormancy during dry periods buffered against environmental filtering ^19,20^.

The transition toward stochastic processes coincided with increased expression of mobility-related genes (Fig. 3d). Key taxa exhibited heightened expression of genes for chemotaxis, flagellar assembly, gliding motility, and pilus systems. For example, Mixococcota expressed flagellar and gliding motility genes, while Actinobacteria, Chloroflexota, and Desulfobacterota activated similar mobility mechanisms. Notably, Pseudomonadota upregulated pilus assembly genes for twitching motility. Enhanced mobility, combined with dormancy strategies, likely homogenized microbial communities while trait-based selection influenced dispersal capabilities ^35,48^.

Metabolite assemblages showed strikingly different patterns, governed by deterministic processes throughout the monsoon cycle, similar to observations in peatland ecosystems ^46^.

Wet periods drove high metabolome variability through increased plant inputs ^49^, secondary metabolite production ^50^, osmolyte release ^51^, and microbial necromass accumulation ^52^.

Concurrently, rainfall-induced runoff and erosion redistributed soil organic matter ^53^, contributing to metabolite homogenization in mid-July.

Dry periods produced a variable metabolome reflecting drought survival strategies. Sugar-like metabolites varied in their role as osmolytes ^54^, while lipid variability suggested membrane remodeling under water-limited conditions ^55^. Abiotic factors, including photodegradation and wind erosion, further shaped organic matter dynamics and metabolome heterogeneity ^56^.

These contrasting assembly patterns show how ecological processes interact to maintain ecosystem function. Stochastic processes maintain microbial community stability during dynamic environmental shifts, while deterministic processes drive metabolite responses, ensuring ecosystem functionality. Together, these mechanisms allow microbiomes to adapt to environmental fluctuations while maintaining their core structure and function.

### Genomic features supporting resilience in arid soils

To understand the genomic basis of microbial stability in arid soils, we next investigated the mechanisms underlying community-level responses. Our genome-centric approach recovered 282 metagenome-assembled genomes (MAGs) (Fig. 4a) of medium to high quality (Supplementary Table 6, Supplementary Fig. 3). These MAGs captured approximately 60% of the sequenced microbial community at the genus level (86% sequence identity), providing a robust foundation for exploring genomic adaptation in arid soils.

**Figure 4.**
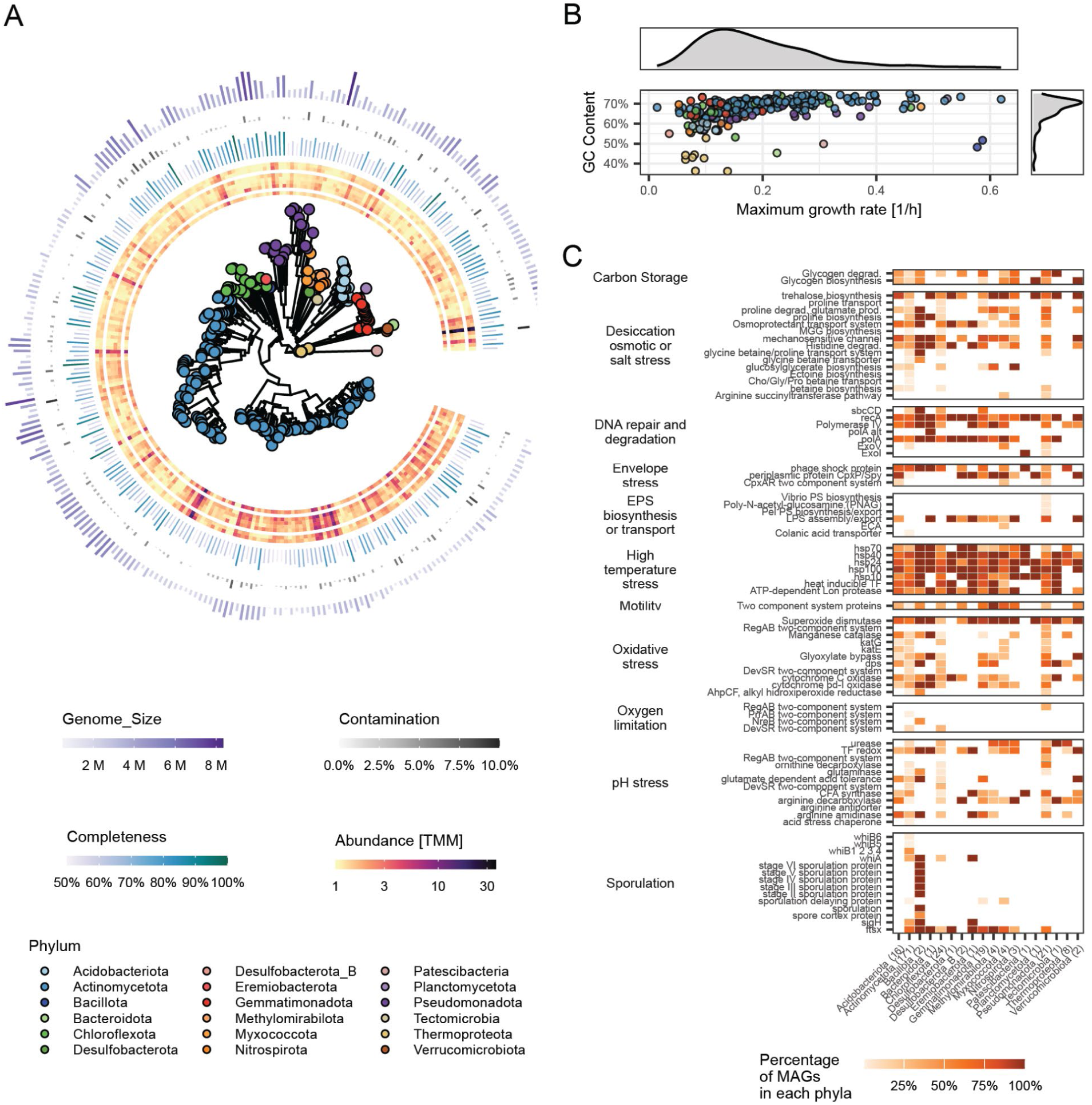
(A) Phylogenetic tree of the 282 recovered metagenome-assembled genomes (MAGs) showing their GC content, genome size, completeness and contamination inferred by CheckM2 as well as their changes in abundance throughout the sampling campaign. (B) Dot plot showing the distribution of the recovered MAGs based on their GC consent and estimated maximum growth rate. (C) Heatmap showing the presence of stress resistance traits among the different taxa of the recovered MAGs.

Phylogenomic analysis revealed unexpected taxonomic breadth, with MAGs spanning 30 bacterial and one archaeal classes across 18 phyla. While this diversity highlights the complex microbial ecology of arid soils, it also suggests high endemism - only one MAG was classified at species level (ANI > 95%), while most (n = 173) were classified at genus level, indicating potentially novel taxa adapted to arid conditions (Supplementary Table 6).

Analysis of carbon fixation pathways revealed how diverse metabolic capabilities are distributed across the community. The Calvin-Benson-Banshan (CBB) cycle appeared in twelve bacterial genomes from Actinomycetota, Desulfobacterota and Pseudomonadota, which encoded complete RuBisCO genes (K01601, K01602) and associated pathway components (Supplementary Fig. 4). While some steps of the reductive citrate (rTCA) cycle were widely distributed, only one MAG from Nitrospirota encoded the key enzyme. Similarly, the 3- Hydroxypropionate/4-Hydroxybutyrate (3HP/4HB) and dicarboxylate/4-hydroxybutyrate (DC/4HB) cycles showed restricted distribution, with key enzymes found only in Thermoproteota, Desulfobacterota and Tectomicrobia.

Nitrogen cycling functions displayed comparable patterns of specialized and redundant distribution. Nitrification capacity was limited to Thermoproteota, specifically the ammonia-oxidizing archaea (AOA) order Nitrososphaerales. In contrast, denitrification and dissimilatory nitrate reduction were broadly distributed across taxa. Sulfate reduction pathways were mostly limited to assimilatory and dissimilatory functions (full details in Supplementary Note 2).

This genomic architecture— featuring both specialized and redundant metabolic functions—highlights how microbial communities maintain critical ecosystem processes under environmental stress. Specialized pathways enable niche-specific adaptations, while functional redundancy across taxa ensures resilience to fluctuating environmental conditions. Together, these features provide a mechanistic understanding of microbial resilience in arid soils.

### Genomic traits reveal convergent and specialized stress adaptations

At the individual level, we investigated how specific traits and stress tolerance mechanisms support microbial survival to the environmental conditions of arid ecosystems. Most recovered MAGs (n = 183) exhibited high GC content and slow predicted growth rates (n = 143, maximum growth rate < 0.2) (Fig. 4b), suggesting a common evolutionary response to harsh conditions ^57^. High GC content likely enhances thermotolerance ^58^, while slow growth rates indicate oligotrophic lifestyles ^59^ aligning with previous findings of GC enrichment in arid conditions ^27^ and resource-driven selection against AT base pairs in nutrient-limited soils ^60^. Notably, Thermoproteota demonstrated an alternative strategy having low GC content, suggesting successful adaptation through genome streamlining ^61^, and efficient nitrogen use ^62^.

Analysis of the gene expression levels of stress tolerance traits (Supplementary Table 7), revealed that protein stability mechanisms were widespread, with most taxa encoding and expressing genes for protection, repair, and degradation of denatured and misfolded proteins (Fig. 4c, Supplementary Fig. 5), reflecting adaptation to high temperatures ^23^. These included various heat shock proteins (HSP70, HSP40, HSP60, HSP10, HSP100, HSP24) critical for protein folding and protection ^63^.

Osmolyte production and transport showed both common and specialized patterns (Fig. 4c, Supplementary Fig. 5). While trehalose and glutamate production was widespread, glycine betaine biosynthesis was restricted to select groups, and bacteria-exclusive ectoine was found only in Actinomycetota. These osmolytes help manage both high salinity and sudden water influx ^63^.

Oxidative stress responses revealed similar diversity in adaptation strategies.

Superoxide dismutase (K04564) showed widespread expression by most MAGs (n = 140) during dry months, particularly a higher expression in Thermoleophilia members (Supplementary Fig. 5). Different taxa employed specialized catalase systems: heme-based catalases (KatG, KatE) in Acidobacteriota, Actinomycetota, Chloroflexota, Myxococcota and Pseudomonadota (KatG n = 41; KatE n = 8), and manganese catalase (K07217) in Thermoproteota, Gemmatimonadoa and Nitrospirota (n = 41) with the highest expression during dry months (Supplementary Fig. 5), suggesting specialized strategies for reactive oxygen species (ROS) management ^64^.

Multifunctional dps proteins (K04047) were present in several phyla (Fig. 4c), providing resistance against oxidative, metal, and thermal stress ^65^, with the highest expression occurring during the dry months for Acidobacteriota, Actinomycetota, Chloroflexota, Gemmatinomonadota, Tectomicrobia and Thermoproteota, and in the wet months for Pseudomonadota (Supplementary Fig. 5). Furthermore, the genes for the glyoxylate bypass, suggested to play a role in oxidative stress ^66^, were found encoded in many taxa and expressed during the dry months in Actinomycetota, Chroloflexota and Tectomicrobia, and during the wet months in Acidobacteriota, Desulfobacterora, Gemmatinomonadota and Pseudomonadota genomes (n = 48) (Supplementary Fig. 5).

Other widely distributed stress tolerance mechanisms included the expression of genes relate to DNA repair as most taxa except for Thermoproteota encoded and expressed genes for the RecA (K03553) protein, which catalyzes one of the central steps of the DNA repair and homologous recombination pathway ^67^ as well as polymerases such as polA (K02335) and polIV (K02346, K04479) that may play a role in double-strand DNA break repair ^68,69^. Other strategies related with dormancy and persistence such as sporulation, exopolysaccharide (EPS) and lipopolysaccharide (LPS) production, and polymeric carbon storage are explored in Supplementary Note 3.

These findings revealed how genomic traits and their modulation throughout the monsoon season support the compositional stability of microbial communities, from an individual standpoint. The combination of shared survival mechanisms and unique adaptations, exemplified by Thermoproteota’s distinct strategies, provides a genomic foundation for community resilience to environmental fluctuations. These insights enhance our ability to predict ecosystem responses to climate change.

### Network restructuring reveals how individual traits shape community responses

To bridge individual adaptations and community-level properties, we analyzed both microbial co-occurrence networks and their metabolic interaction potential. Using a random matrix theory (RMT)-based approach ^70^, we found that increased precipitation dramatically altered network architecture ^71,72^. During dry periods, networks were larger and more modular (higher modularity, transitivity, and node/edge numbers, lower average path distance), whereas wet conditions produced smaller, more hierarchical networks (higher degree centralization, betweenness centralization, and stress centrality) (Fig. 5a and Supplementary Table 8). The dramatic shift in network architecture between dry and wet conditions represents more than structural reorganization - it reveals fundamental changes in community interaction strategies. During dry periods, the larger, more modular networks reflect a community-wide strategy for resource conservation. Each module functions as a metabolically interdependent unit, allowing efficient resource cycling within tightly connected groups. This organization stands in stark contrast to conventional understanding of drought responses, where networks typically become fragmented and less connected.

**Figure 5.**
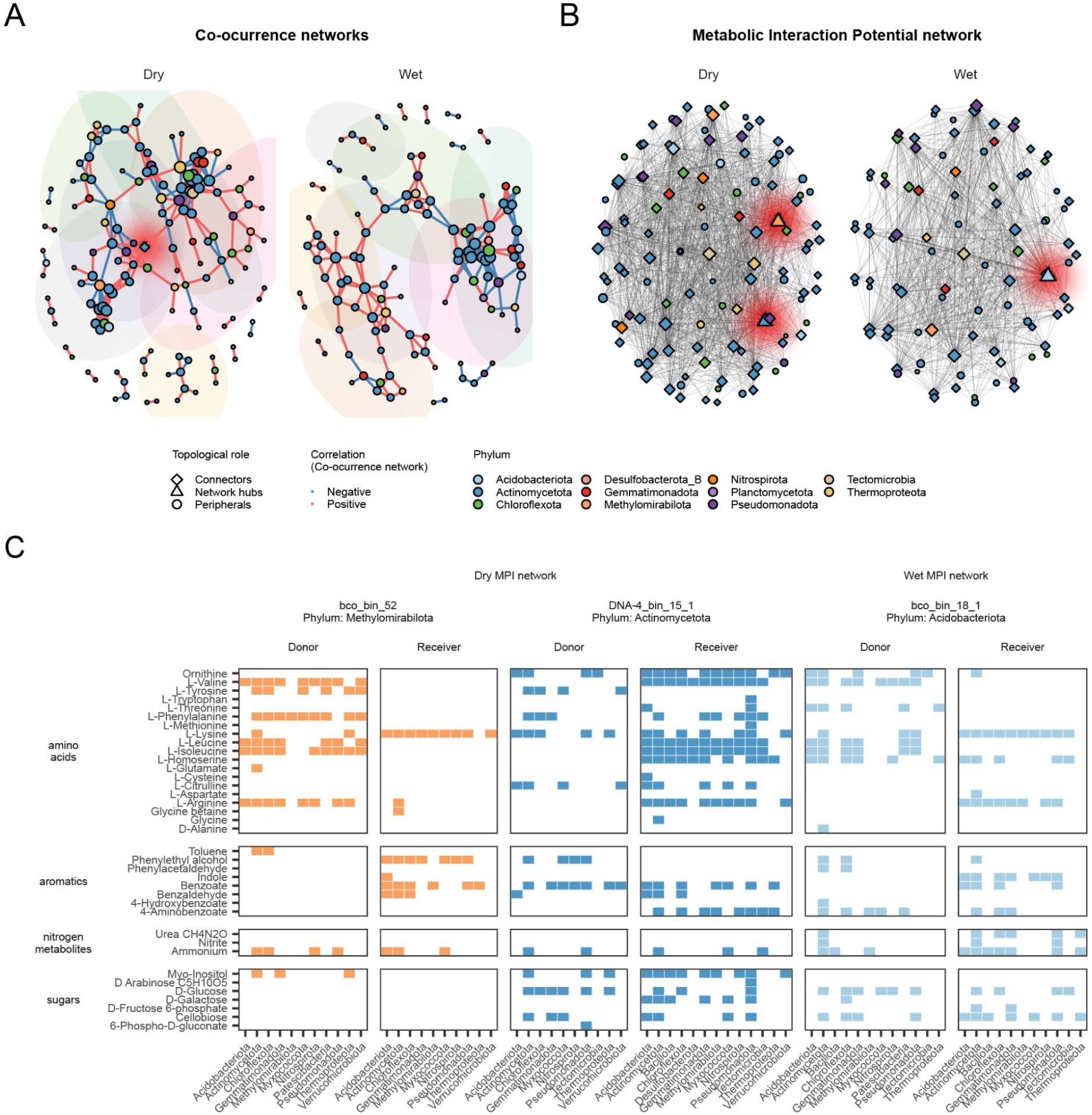
Microbial networks. (A) Co-occurrence networks based on MAGs abundances (TMM) inferred with CoverM for the dry (May and October) and wet (July and August) months. Nodes represent MAGs and are colored based on their Phylum. Node size indicates the Degree (number of connections) of each node. Colored ellipses indicate network modules with more than 6 nodes. (B) Social networks based on the metabolic interaction potential (MIP) score calculated with SMETANA. Only genome pairs with MIP >= 5 were considered for the network. Nodes represent MAGs and are colored based on their Phylum. Node size indicates the Degree (number of connections) of each node. (C) Heatmap showing metabolite donation and uptake patterns between Network Hub microbes and members of other phyla.

Our findings overturn traditional expectations about water availability’s impact on microbial networks. Unlike other ecosystems where wet conditions promote higher interconnectedness ^73^, dry conditions in our system fostered tighter community organization through modular, interconnected network structures. This tight organization likely reflects the need for efficient resource utilization ^74^, where interdependence enhances environmental resilience. Conversely, wet conditions relaxed these tight resource constraints, allowing for more hierarchical organization with diffuse, opportunistic interactions ^73,75^. These changes in structure suggest microbial interactions and responses play a role in community-level resilience. The higher centrality of wet networks suggests wet environments favor dominant species that quickly capitalize on increased nutrients ^76^, while osmotic stress forces other microbes to regulate metabolism more intensively ^28^. This network restructuring aligns with increased selection processes at monsoon onset (Fig. 3b), as water simultaneously drives community structure and metabolic activity changes ^77,78^, while enhancing niche connectivity through dispersal mechanisms ^4^.

Network analysis identified key species bridging individual traits and community function.

In dry conditions, an Actinobacteria MAG served as a network connector, similar to findings from other deserts including Namib ^79^, Atacama ^80^, and Sahara ^81^. However, this MAG’s high abundance challenges the assumption that desert community stability relies on rare taxa ^38^, suggesting abundant species can be central to ecosystem function.

To understand how network changes reflect metabolic adaptations, we analyzed metabolic interaction potential (MIP) using SMETANA ^82^. Dry networks showed more extensive metabolic interactions, indicating increased interdependence and active metabolite exchange ^74^. Moreover, this analysis identified additional potential keystone MAGs (network hubs): Methylomirabilota and Actinomycetota in dry conditions, and Acidobacteriota during wet periods (Fig. 5b). The Methylomirabilota keystone is particularly significant given its roles in greenhouse gas cycling ^83^, soil processes ^84^, and vegetation interactions ^85^. SMETANA analysis in detailed mode (Fig. 5c), revealed that this MAG primarily donates amino acids, as previously reported ^86^, while the Actinomycetota hub absorbs them demonstrating how individual metabolic capabilities shape community-level interactions and eventually resilience through energy-expensive amino acid exchange ^87^. Analysis of metatranscriptomics data confirmed some of these results, as complete biosynthetic pathways for valine, leucine and isoleucine were found to be present in the Methylomirabilota genome (Supplementary Fig. 6a), which were expressed through the monsoon season (Supplementary Fig. 6b). Additionally, expression of amino acid transporters was detected in a wide range of phyla (Supplementary Fig. 6c) indicating the ability of these microbes to retrieve amino acids from the environment. These insights into network restructuring demonstrate how community stability and metabolic responses are interconnected through dynamic organizational changes. This coordination between resilience levels may be crucial for maintaining ecosystem function amid climate change-related precipitation alterations.

### Strategic gene regulation by Thermoproteota maintains critical ecosystem processes

To understand how individual taxa maintain specific functions within changing networks, we examined Thermoproteota as a model organism. We selected this phylum because: (1) eight MAGs from the Nitrososphaeraceae family represented the only aerobic ammonia oxidizers in our community, (2) nitrogen availability critically influences arid ecosystem function ^88,89^ and (3) Thermoproteota’s genomic features and regulatory patterns provide a unique model for understanding how individual adaptations support community stability.

Using genome-resolved metatranscriptomics, we tracked Thermoproteota’s gene regulation throughout the monsoon season (Supplementary Table 9). The ammonia oxidation genes amoABC (K10944, K10955, K10946) showed unexpected expression patterns, peaking during dry periods (Fig. 6a), with amoC (K10946) showing highest expression ^90^. This counterintuitive pattern reveals a sophisticated adaptation strategy. During dry periods, when most microbes reduce activity, Thermoproteota maintains high nitrification rates, likely capitalizing on reduced competition for limited nitrogen resources. This strategy is supported by unique stress tolerance mechanisms, including specialized protein repair systems, manganese-based catalases for ROS management, and efficient nitrogen utilization enabled by genome streamlining. Together, these adaptations allow Thermoproteota to maintain critical ecosystem functions even when other taxa become dormant, consistent with its crucial role in hyperarid soils ^91^.

**Figure 6.**
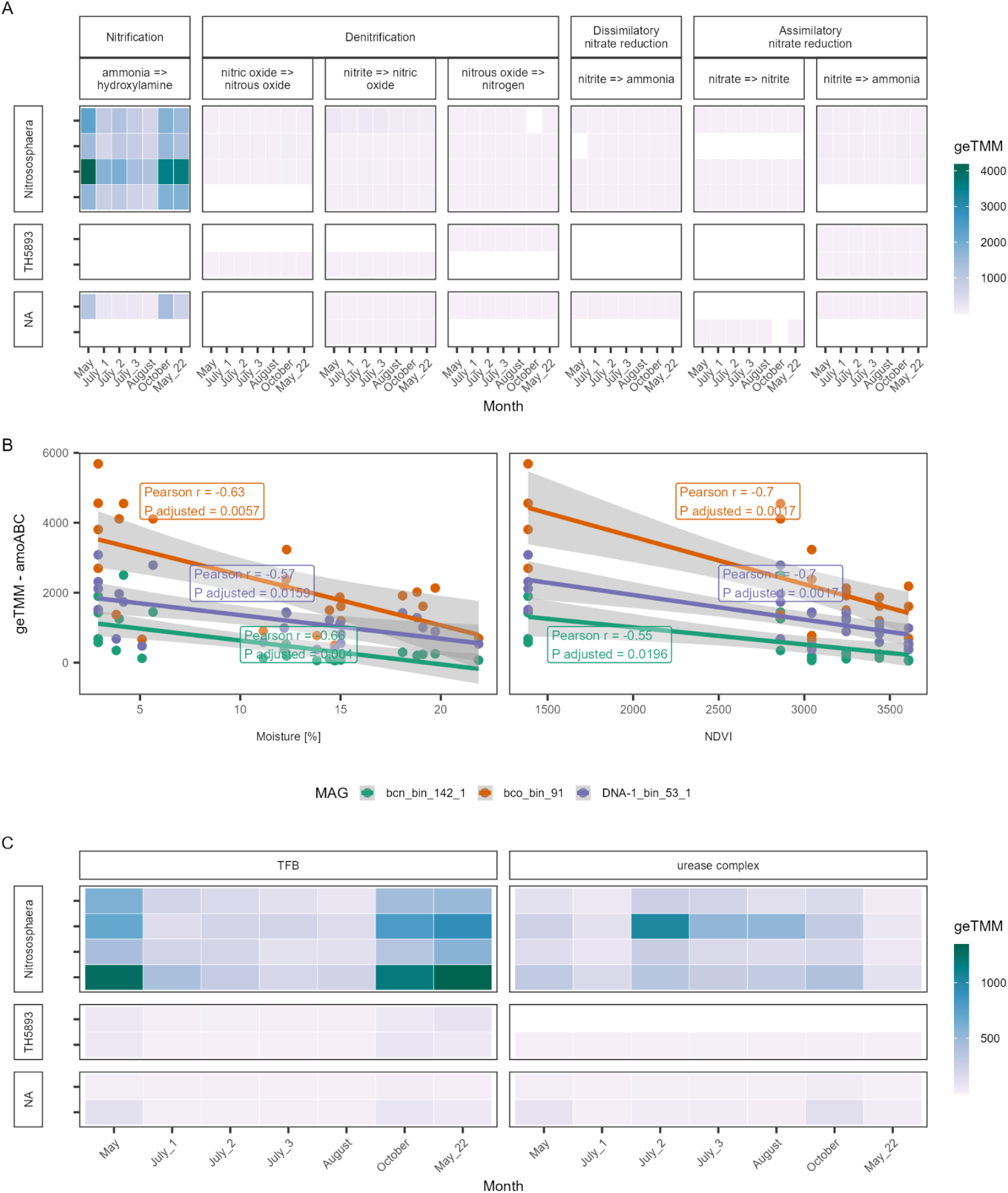
(A) (A) Heatmap of nitrogen metabolism gene expression (geTMM) across Thermoproteota genomes. (B) Dotplot showing correlations between amoABC gene complex expression and environmental variables (soil moisture and NDVI). (C) Heatmap of urease complex and transport gene expression (geTMM).

Environmental correlations further highlight how individual adaptations respond to community-level conditions. A strong negative correlation between amoABC expression and both soil moisture and NDVI (Fig. 6b, Supplementary Table 1) suggests that Thermoproteota sustains high nitrification activity under the driest conditions, potentially outcompeting other microbes for limited nitrogen. Moreover, four Thermoproteota MAGs showed high expression of the urease gene cluster (ureABCDEFG) during wet periods (Fig. 6c), indicating these taxa can metabolize both ammonia and urea as energy sources ^92^. This versatility exemplifies how individual-level adaptations enable persistence across variable conditions, providing significant ecological advantages in nutrient-limited soils.

The absence of ammonia-oxidizing bacteria (AOB) in our system further highlights how Thermoproteota’s strategy of regulated persistence enables their dominance of nitrification processes. Analysis of the taxonomic profiles of other arid soil datasets showcase the widespread presence of Thermoproteota in arid systems worldwide (Supplementary Fig. 2h). This global distribution underscores their importance in stabilizing nitrogen fluxes, a critical ecosystem function in arid environments that are particularly vulnerable to climate change-induced disruptions ^89^, aligning with previous reports of archaeal dominance in arid soils ^89^ and their broader distribution across oligotrophic environments ^93,94^. Their success demonstrates how individual-level expression modulation contributes to community-level resilience by maintaining essential ecosystem processes despite environmental stress (Supplementary Note 4).

The success of this strategy extends beyond individual survival to shape community-level functions in several ways. First, Thermoproteota’s maintained activity during dry periods provides a consistent nitrogen supply that supports community stability. Second, its network position shifts from peripheral during wet periods to central during dry periods, demonstrating how individual adaptations influence community organization. Third, its regulated persistence exemplifies how specialized taxa can maintain essential ecosystem processes despite environmental stress, contributing to overall community resilience.

These findings expand our understanding of Thermoproteota’s role in arid ecosystem nitrogen cycling ^38^ and demonstrate how specialized adaptations maintain critical functions supporting community stability. This coordination between individual regulation and community-level processes provides insights into how arid ecosystems may persist amid increasing climate variability.

## DISCUSSION

In this study, we applied an integrated time-resolved multiomics approach to better understand how microbial communities can respond to dramatic environmental transitions. Our findings reveal the interplay between community-level adaptations and individual stress responses, providing a comprehensive view of how microbial networks reorganize dynamically to sustain ecosystem functions under fluctuating conditions. Using soils from the Sonoran Desert as a model system, characterized by the highly variable environmental conditions of the monsoon season, we identified three interconnected mechanisms underpinning microbial resilience: (1) stochastic processes that maintain community stability, (2) dynamic network reorganization, and (3) coordinated individual stress responses. Additionally, we identified Thermoproteota as a key taxon in arid soils, highlighting its potential as a model organism for understanding adaptation to aridity.

### Microbial Network Responses to Environmental Fluctuations

Our findings challenge the prevailing view that drought predominantly disrupts microbial networks ^72,95,96^. Instead, we observed enhanced microbial network connectivity under dry conditions, aligning with evidence that stress can foster tighter microbial associations ^97^. This tightly organized community structure became more diffuse during wet conditions, suggesting dynamic reorganization based on resource availability (Figure 5A, Supplementary Table 8).

These results may reflect unique adaptations of microbial communities in bare soils, where plant buffering effects are limited ^24,95^.

The observed increase in network connectivity under drought conditions supports the stress gradient hypothesis ^98–100^. During dry periods, certain taxa, such as Thermoproteota and Actinomycetota, likely act as hubs, facilitating functional complementarity and efficient resource sharing among community members. Conversely, wet conditions promoted larger, more centralized networks with increased negative correlations, indicating the rise of dominant microbes and heightened competition ^75,100^. This dynamic network restructuring is crucial for maintaining microbial diversity and ecosystem function, as evidenced by stable community composition across monsoon cycles and the dominance of stochastic assembly processes (Figure 3C). These patterns highlight how microbial communities balance stability and adaptability in fluctuating environments.

### Network Reorganization Bridges Community Stability and Individual Adaptations

The link between network reorganization and resilience underscores the importance of coordination at multiple biological levels ^6^. Tight networks during dry periods optimize resource exchange and foster interdependencies, enhancing community stability under stress ^82,101^. This is supported by correlations between network modularity and the expression of stress-response genes. In contrast, diffuse networks during wet periods allow metabolic flexibility and opportunistic interactions, enabling communities to rapidly exploit resource pulses while maintaining core functions ^12^.

Despite inherent limitations in metagenomics-derived network analyses—such as the inability to discern the precise nature of correlations, which may arise from indirect interactions or shared niches ^102^—our approach demonstrates the utility of network analysis for inferring ecological functions ^101,103,104^. Here, we go further ahead and provide some evidence on how the sophistication of this network-mediated coordination emerges through metabolic interactions.

For example, the prominent role of Thermoproteota during dry conditions (Fig. 5A) highlights its contribution to nitrogen cycling, a critical process for maintaining ecosystem function. Its widespread presence in arid environments globally (Supplementary Fig. 2g) further emphasizes its ecological significance as both an indicator of resilience and a potential biotechnological target. Meanwhile, Methylomirabilota, Actinomycetota and Acidobacteriota maintained active amino acid exchange with other community members (Fig. 5C), supported by sustained expression of biosynthetic pathways and transporters (Supplementary Fig. 6). These coordinated interactions create reciprocal dependencies optimizing resource utilization, forming feedback loops where individual metabolic activities shape community structure, while community organization influences individual responses ^105^.

### Organic Matter Profiles Reflect Coordinated Stress Responses

The analysis of organic matter profiles reveals deterministic changes driven by both community-level metabolic adjustments and individual stress responses ^46^. Unlike plant- dominated ecosystems ^36,37^, the prevalence of lipid- and protein-like compounds in these soils underscores the strong microbial influence on soil organic matter composition ^106^. During dry periods, the accumulation of osmolytes and stress-related compounds reflects individual microbial adaptations ^107^, while shifts in metabolite profiles during wet periods indicate community-level metabolic reorganization. These changes mirror network restructuring, as metabolite exchange relationships transition between tight and diffuse network arrangements, further illustrating the integration of microbial activities in maintaining ecosystem resilience.

### Individual-Level Survival Strategies and Their Role in Resilience

Our analysis uncovered shared and specialized survival strategies among taxa, highlighting strong selective pressures for traits that support both individual survival and community function ^27^. For instance, Actinomycetota employed multiple adaptations including high GC and production of chaperone proteins to to provide thermostability and protect cellular machinery ^108^, spore formation to enable dormancy ^109^, osmolyte production to overcome water stress ^110^, and versatile metabolic strategies to allow resource optimization ^111^. Similarly, the Thermoproteota maintained regulated function through sophisticated stress response mechanisms including chaperones, DNA-repair proteins and the widespread production of stress-related proteins like katG, katE, dps, and superoxide dismutases to overcome stressful conditions while maintaining regulated function in the nitrification pathway (Fig. 6a, Supplementary Fig. 4). The consistency of stress response mechanisms across diverse taxa suggests strong selective pressure for maintaining both individual survival and community function ^112^. The widespread distribution of heat shock proteins, osmolyte production pathways, and oxidative stress responses indicates that these mechanisms represent essential adaptations to arid conditions ^19,96^. Together, these diverse survival strategies underscore the critical role of functional redundancy and metabolic versatility in maintaining community stability, demonstrating how alternative strategies can achieve similar community-level outcomes ^19,20^.

Such individual-level adaptations form the building blocks of network-mediated resilience, enabling the community to buffer environmental perturbations and sustain key ecosystem processes.

### Implications for Microbial Resilience and Climate Change

Our findings provide a framework for understanding microbial resilience in arid systems. Specifically, stochastic processes ensure stable community structures, while dynamic network reorganization and individual stress responses enable adaptati*o*n to extreme environmental conditions (Figure 7). The pivotal role of Thermoproteota in nitrogen cycling highlights its potential as both an ecological indicator and a target for biotechnological interventions aimed at enhancing soil resilience. Furthermore, the observed prevalence of shared stress response mechanisms and network reorganization patterns may suggest that the principles of network-mediated resilience revealed in our study likely extend across arid ecosystems worldwide and potentially non-arid ecosystems ^113^.

**Figure 7.**
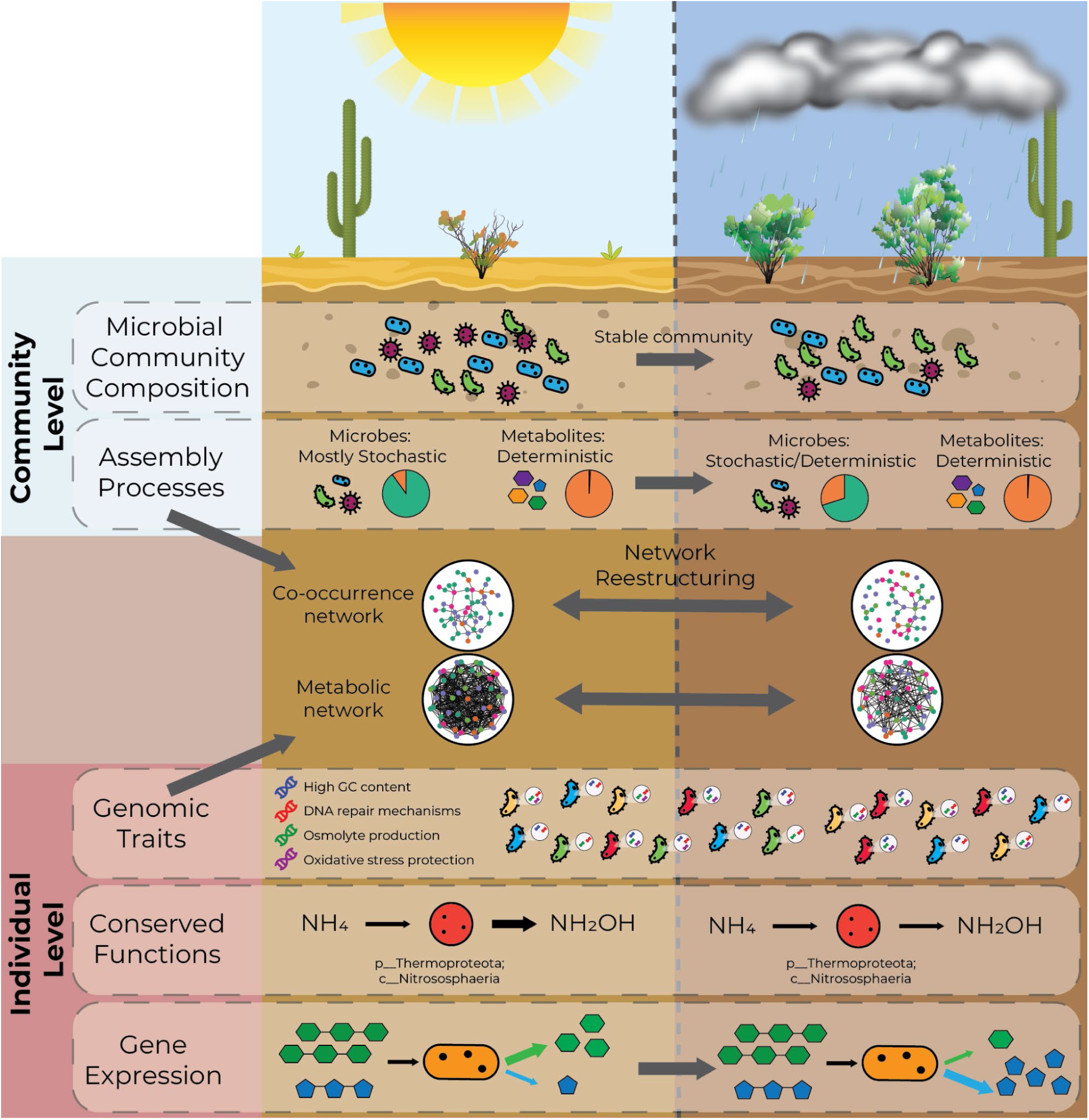
Proposed dual-level resilience framework showing microbial responses to monsoon fluctuations in arid ecosystems. Resilience emerges from interactions between individual-level mechanisms (genomic traits, metabolic functions, gene regulation) and community-level processes (composition, assembly). These interactions drive network restructuring, enabling resource sharing and protection during dry periods, and resource exploitation during wet periods. Individual-level adaptations include high GC content, DNA repair mechanisms, and osmolyte production, while community-level responses involve both co-occurrence and metabolic interaction networks.

While these multi-level resilience mechanisms currently enable functional persistence through monsoon cycles, they also suggest vulnerabilities to future climate change. Increased aridity, prolonged droughts, or disruptions to stochastic processes may exceed the adaptive capacity of these specialized microbial communities, leading to cascading effects and destabilization ^28,73,114,115^. To safeguard critical ecosystem functions, future efforts should prioritize promoting functional redundancy within microbial communities, conserving microbial diversity hotspots, and monitoring network reorganization thresholds. Additionally, integrating predictive modeling with multi-omics datasets could facilitate early detection of resilience breakdowns under changing climate scenarios. Tracking key taxa such as Thermoproteota will be essential to these efforts, providing critical data on how microbial communities navigate environmental extremes and informing strategies to safeguard ecosystem functionality in a rapidly changing world.

## MATERIALS AND METHODS

### Soil collection

Soil samples were collected at the Saguaro National Park West, Tucson, Arizona (32° 15 ’05.4”N, 111° 09’ 40.2”W), using ethanol-sterilized shovels, at 10-15 cm depth. Sample collection occurred across 9 time points spanning pre-monsoon (May 2021, 2022), monsoon (June-August 2021), and post-monsoon (September-October 2021) seasons, with 4 biological replicates per time point (total n = 36). Samples were transported on ice to the laboratory, and sequentially sieved through 4000 µm and 2000 µm Fieldmaster® sieves (Science First, Yulee, FL). Samples were stored at -80°C until analysis.

The sampling site is located within the Sonoran Desert in the Southwestern USA. A region that experiences extreme temperatures exceeding 40°C, with maximum temperatures reaching 48°C ^116^. It has an annual precipitation average of 200 mm (1980-2016), with regional variations across the desert landscape ^117^. The area’s distinctive bimodal precipitation pattern includes winter frontal system rainfall and summer monsoon storms ^118^, with a characteristic dry period in late spring.

### Soil physicochemical properties

For soil moisture, around 5 g of soil were placed in 15 ml Eppendorf tubes (3 replicates per sample) and lyophilized on a FreeZone 2.5 Liter Benchtop Freeze Dryer (Labconco, Kansas City, MO) for 72 hours. Moisture content (%) was calculated as: [(initial weight - lyophilized weight)/initial weight] × 100. For pH measurements, 2 g of soil were mixed with 6 ml double-distilled water, vortexed for 15 minutes, centrifuged for 3 minutes, and incubated at room temperature for 30 minutes. pH was measured twice per sample using a benchtop Orion Star A211 pH meter (Thermo Scientific, Waltham, MA).

Total carbon (TC) and nitrogen (TN) were measured for all samples (n=36) using a ECS 4010 Elemental combustion analyzer (Costech Analytical Technologies, Valencia, California) by the Arizona Laboratory for Emerging Contaminants (ALEC) at the University of Arizona. Major cations (K, Ca, Na, Mg) and trace metals (V, Cr, Fe, Co, As, Se) were quantified using an Agilent 7700 inductively coupled plasma mass spectrometer (ICP-MS) (Agilent, Santa Clara, CA) as described before ^119^ with the exception that the US EPA Method 3051A was used for soil acid digestion. All measurements are available in Supplementary Table 1.

Daily precipitation data, soil and air temperature measurements were retrieved from the U.S. Climate Reference Network Data website (https://www.ncei.noaa.gov/pub/data/uscrn/). Normalized difference vegetation index values were retrieved from the Terra Moderate Resolution Imaging Spectroradiometer (MODIS) Vegetation Indices (MOD13Q1) Version 6.1 ^120^ dataset.

For multivariate analysis of the environmental variables, they were first standardized with the vegan package. Standardized values were used to produce a hierarchical clustering of the samples using Euclidean distances. Additionally, a principal component analysis (PCA) was also performed with this data to better visualize the differences between samples.

### FTICR-MS sample preparation and data preprocessing

Ultra-high resolution soil organic matter profiling was conducted using Direct Infusion Fourier Transform Ion Cyclotron Resonance Mass Spectrometry (FTICR-MS). Soil metabolites were extracted through sequential water and methanol extraction. A total of 2.5 g of each 2021 soil sample (n = 32) was mixed with 5 ml of dd 15 ml Eppendorf tubes with 5 ml of ddH2O, followed by two rounds of vortexing and 2-hour sonication. Samples were then centrifuged at 5000 rpm for 5 minutes. The supernatant was collected and stored at -20°C. The remaining soil mix underwent two rounds extra of extraction with the same protocol using 5 ml of HPLC-grade methanol instead of ddH2O. Combined water and methanol extracts were purified using solid phase extraction (SPE) to remove contaminants ^121^. Extracts were eluted using 1.5 mL of HPLC methanol, and the final purified extracts were shipped to the Pacific Northwest National Laboratory (PNNL) for mass spectrometry analysis.

High-resolution mass spectra data was collected with a 12-Tesla Bruker SolariX FTICR mass spectrometer (Bruker, SolariX, Billerica, MA) located at the Environmental Molecular Sciences Laboratory at PNNL in Richland, WA. Samples were directly infused into the instrument using a custom automated direct infusion cart that performed two offline blanks between each sample. The FTICR-MS was outfitted with a standard electrospray ionization (ESI) source, and data was acquired in negative and positive mode with the needle voltage set to +4.2kV and -4.2kV respectively. Data was collected at 4MW, and the ion accumulation time (IAT) was optimized for each polarity. In negative mode the IAT was set to 0.08 sec and data was measured from 98.28 m/z to 2000 m/z with a resolution of 300K at 333.1118 m/z. In positive mode the IAT was set to 0.08 sec and the data were measured from 147.42 m/z to 1400 m/z with a resolution of 380K at 385.1176 m/z. One hundred forty-four scans were co-added for each sample and internally calibrated using OM homologous series separated by 14 Da (–CH2 groups). The mass measurement accuracy was typically within 1 ppm for singly charged ions across a broad m/z range (100 m/z - 900 m/z). Bruker Data Analysis (version 4.2) was used to convert raw spectra to a list of m/z values by applying FTMS peak picker module with a signal-to-noise ratio (S/N) threshold of 7 and absolute intensity threshold of 100.

Chemical formulae were then assigned using Formularity ^122^, with parameter: S/N >7, and mass measurement error < 0.5 ppm, allowing only for the presence of C, H, O, N, S and P. We present results from positive mode analysis only, which revealed the strongest shifts in metabolite composition.

Thermodynamic indexes for the FTICR molecular formulae, chemodiversity indexes and putative biochemical transformations were calculated using MetaboDirect (version 1.0.7) ^123^ keeping only peaks that were present in at least 4 samples and with the sum normalization method. The number of nitrogen and sulfur containing transformations were inferred based on the assigned transformation names (Supplementary Table 2).

### Metabolite dendrogram construction

To analyze monsoon-mediated shifts in metabolite assembly processes in arid soils, we applied ecological null modeling methods originally developed for microbial data ^43,44^, and then adapted for the analysis of metabolites ^45,46^. Metabolites were categorized into three groups based on molecular class assignments ^124^: group 1: “lipid-like”, group 2: carbohydrate-, protein- and amino sugar- like compounds referred to as “sugars” and group 3: tannin-, lignin- and condensed hydrocarbon-like compounds referred to as “TLC”.

Relational dendrograms were constructed for both bulk metabolite data and the three metabolite pools using three approaches 1) metabolite molecular characteristics (MCD), 2) metabolite potential biochemical transformations (TD), and 3) metabolite transformation-weighted characteristics (TWCD) ^45^. Dendrogram construction utilized binary presence/absence values to eliminate charge competition bias in abundance estimates ^45^.

### DNA extraction

DNA was extracted from 250 mg soil samples using the DNeasy PowerSoil Pro Kit (Qiagen, Hilden, Germany) according to manufacturer protocols. DNA quantification was performed using the Qubit DNA high sensitivity kit (ThermoFisher Scientific, Waltham, MA). Separate DNA extractions were conducted for 16S amplicon sequencing and shotgun metagenomics. The University of Arizona Genetics Core (RRID: SCR_012429) performed all library preparation and sequencing data generation.

### 16S rRNA amplicon sequencing

Amplicon sequencing was conducted on samples collected from May to October 2021 (n = 32). The V3-V4 regions of the 16S rRNA gene were amplified using previously designed primers ^125^ (Forward: 5’-CCTACGGGNGGCWGCAG-3’ and Reverse: 5’- GACTACHVGGGTATCTAATCC-3’), with Illumina™ overhang adaptors at their 5′ ends. Amplicon amplification was performed using KAPA HiFi HotStart Ready Mix (Roche Sequencing Solutions, Pleasanton, CA) following manufacturer protocols. A secondary PCR utilized Nextera XT v2 Index kits sets A-D (Illumina, San Diego, CA). Amplicon purification employed HighPrep PCR magnetic beads (MagBio, Gaithersburg, MD) at 0.8X and 1.2X volumes for first and second PCR products, respectively. Libraries were quantified using QuantiFlor dsDNA System (Promega, Madison, WI) on a BioTek FLX 800 plate reader (BioTek, Charlotte, VT). Size determination of pooled amplicons used an AATI Fragment Analyzer with HS NGS Fragment kit (Agilent, Santa Clara, CA). Pools were diluted to <20nM and quantified using a Roche LightCycler 480-II with KAPA Illumina Library Quantification Kit (Roche Sequencing Solutions, Pleasanton, CA). Sequencing was performed on an Illumina MiSeq platform using the v3-600 cycle kit (Illumina, San Diego, CA), generating 250 bp paired end reads.

### Metagenomics sequencing

Metagenomic libraries were prepared from samples collected in 2021 during May (Sampling time point 1; n = 4), July (Sampling time points 3, 4 and 5; n = 12), August (Sampling time point 6; n = 4), October (Sampling time point 8; n = 4), and May 2022 (Sampling time point 9; n = 4). Quality control involved genomic DNA (gDNA) quantification using QuantiFlor dsDNA System on a BioTek FLX 800 plate reader, and size and integrity assessment using AATI Fragment Analyzer with HS NGS Genomic DNA kit. Genomic DNA (350-550 ng) was sheared to 200 bp using standard conditions on an S2 sonicator (Covaris, Woburn, MA). Library preparation utilized NEBNext Ultra II DNA Library Prep Kit for Illumina (New England Biolabs, Ipswich, MA) with NEBNext Multiplex Oligos for Illumina UDI primer pairs (New England Biolabs, Ipswich, MA), employing 3 PCR enrichment cycles. Libraries were purified using 0.8X volume HighPrep PCR magnetic beads, quantified with QuantiFlor dsDNA System on a BioTek FLX 800 plate reader, and sized using AATI Fragment Analyzer with HS NGS Fragment kit.

Size-adjusted libraries were quantified using Roche LightCycler 480-II with KAPA Illumina Library Quantification Kit before equal pooling. Initial metagenomic sequencing generated 2x150 bp paired-end reads using one full lane of a NovaSeq 6000 S4 flow cell v1.5 (300 cycles) on the NovaSeq 6000 platform (Illumina, San Diego, CA). A second sequencing round was performed for six selected samples (one per time point) using another full lane of a NovaSeq 6000 S4 flow cell v1.5 (300 cycles).

### RNA extraction and sequencing

Metatranscriptomic sequencing was conducted on the same samples used for metagenomic analysis (n = 28). RNA was extracted from 2 g soil samples using RNeasy PowerSoil Total RNA kit (Qiagen, Hilden, Germany) following manufacturer protocols. DNA removal was performed using DNase I (New England Biolabs, Ipswich, MA), followed by additional purification using RNA Clean Kit T2030 (New England Biolabs, Ipswich, MA). Initial RNA quantification employed Qubit RNA high sensitivity kit (ThermoFisher Scientific, Waltham, MA), with RNA integrity assessed via 2100 Bioanalyzer (Agilent, Santa Clara, CA) before submission to the University of Arizona Genetics Core (RRID: SCR_012429) for library preparation and sequencing.

Prior to library preparation, secondary RNA quantification was performed using Qubit Fluorometer 2.0 with Qubit Broad Range RNA Quantification Kit (ThermoFisher Scientific, Waltham, MA), and RNA quality was evaluated using AATI Fragment Analyzer with 15 nt RNA Kit. RNA ribodepletion was conducted on 50-500 ng input RNA using NEBnext rRNA Depletion Kit (Bacteria) (New England Biolabs, Ipswich, MA). Libraries were prepared using NEBNext Ultra II Directional RNA Library Prep Kit for Illumina (New England Biolabs, Ipswich, MA) and NEBnext Multiplex UDI Primer Pairs for Illumina (New England Biolabs, Ipswich, MA), with 9-14 PCR enrichment cycles. Double purification of libraries used 0.8X volume HighPrep PCR magnetic beads. RNA library quantification and sequencing followed the same protocols described for DNA libraries.

### Comparison with other datasets

Metagenomics read signatures were compared with shotgun metagenomics reads available in public databases from other arid ecosystems across the world including the Negev Desert (PRJEB36534) ^19^ and (PRJNA657906) ^126^, Sinai Desert (PRJNA810925) ^127^, Xilingol League (PRJNA916612) ^128^, Mojave Desert (PRJNA620793) ^129^, Cuatro Cienagas (PRJNA857298) ^130^, as well as our same study ecosystem, the Sonoran Desert (PRJNA664514) ^131^, using MASH distances ^132^. Microbial taxonomic profiles of all datasets were inferred with SingleM (v0.18.2) ^133^.

### Microbial community profiling and analysis

Microbial community profiles based on amplicon sequence variants (ASVs) were generated using 16S rRNA gene amplicon sequencing reads with the DADA2 (v1.26) pipeline ^134^. Briefly, adapter trimming was performed using cutadapt ^135^, trimmed reads were then filtered and quality trimmed to a minimum length of 50 and maximum expected error of 2. Error rates were learned directly from processed reads using DADA2 sample inference algorithm to infer sequence variants. Fully denoised sequences were generated by merging ASVs from forward and reverse reads followed by chimeric sequence removal. Taxonomy was assigned using a naive Bayesian classifier ^136^ with Genome Taxonomy Database (GTDB) version r220 ^137^.

To account for 16S rRNA gene copy number variation effects on community analysis ^138^, metagenomic-based taxonomic profiling was performed using SingleM (v0.18.2) ^133^ on metagenomic raw reads following tool authors’ recommendations. Alpha diversity indexes were calculated as the average value of the indexes obtained from each marker gene. For other downstream analyses we used the *rplP* gene (encoding ribosomal protein L16 L10e), due to its effectiveness in distinguishing between closely and distantly related genomes ^139^.

Statistical analyses were conducted in R (v4.3.2) ^140^. Both datasets were rarefied to 25,000 reads for alpha and beta diversity calculations, while other analyses used unrarefied datasets. Observed and estimated richness (Shannon diversity index and Chao1) were calculated with the *phyloseq* R package ^141^, while beta diversity analysis was done with Bray Curtis distances calculated with the *vegan* package ^142^. Principal coordinates analysis (PCoA) was performed using *phyloseq*, with PERMANOVA testing for differences between sampling time points.

For phylogenetic analysis of 16S rRNA data, multiple sequence alignment was done using the DECIPHER package (version 2.26.0) ^143^. A maximum-likelihood phylogenetic tree was constructed using FastTree (version 2.1.1) ^144^, with a generalized time-reversible model and gamma option for branch length rescaling and Gamma20-based likelihood computation. The tree was midpoint-rooted using the phytools package (version 2.3.0) ^145^.

### Metagenome processing and assembly

Sequencing generated 44-180 million read pairs during the first round and 416-588 million read pairs during the second round per sample. Read quality assessment was performed using FastQC (v0.11.9) ^146^. Raw reads underwent preprocessing using the ”RQCFilter” pipeline (BBtools version 38.90) ^147^. Cleaned reads were assembled with MEGAHIT (v1.2.9) ^148^ with the ‘--presets meta-largè option as recommended for high-biodiversity soil metagenomes. To optimize metagenome assembled genome (MAG) recovery, eight metagenome assemblies were generated using two strategies: 1) Co-assembly of samples from identical sampling points (6 assemblies) and 2) Co-assembly of samples from the same sequencing run (2 assemblies). Assemblies were then filtered to retain only contigs larger than 2000 bp. In total we retrieved between 0.3 and 1.5 million contigs from each assembly with a N50 between 3200 bp to 3500 bp. Read mapping to the filtered assemblies using CoverM (version 0.7.0) ^149^, with flags -p minimap2-sr -m trimmed_mean --min-read-percent-identity 0.95 --min-read-aligned-percent 0.75, indicated they accounted for 29.74% to 42.89% of the reads of each sample.

### Recovery of metagenome assembled genomes

For MAG recovery, filtered contigs of each assembly were binned separately using CONCOCT (v1.0.0) ^150^, MetaBAT2 (v2.12.1) ^151^, and MaxBin2 (v2.2.6) ^152^ with default options. The resulting bin sets were refined using the bin refinement module from metaWRAP (v1.3.0) ^153^. Refined bins were manually curated using anvi’o (version 8) ^154^. Bin quality and completeness were checked using CheckM2 (v1.0.2) ^155^. Only bins with more than 50% completeness and less than 10% contamination (medium quality draft genomes) were retained and used for the analysis resulting in a set of 661 bins. Bins generated from the different assemblies were dereplicated at 95% ANI similarity (species-level ^156^) using galah (v0.4.0) ^157^ resulting in 282 dereplicated MAGs. Read mapping using CoverM, with the same parameters as before, revealed that the recovered MAGs accounted for 3.40% to 9.11% of the reads from each sample, comparable to other metagenomics studies of soil microbial communities (2.7%- 22.4%) ^158^. To determine how much of the microbial communities was represented by the recovered MAGs, we compared the taxonomic profiles of the reads, the assemblies and the recovered MAGs using SingleM appraise at 86% sequence identity (genus level) and the – imperfect flag.

Dereplicated bins were taxonomically classified using the Genome Database Taxonomy toolkit (GTDB-tk) (v2.4.0) ^159^ and the R09-RS220 database. Phylogenetic trees for bacterial and archaeal species were built separately using the de-novo workflow of GTDB-tk based on 120 marker genes for bacteria and 53 marker genes for archaea. Trees were rooted using the p Patescibacteria and p Altiarcheota for the bacteria and the archaeal tree respectively. Visualization of the trees was performed using the ggtree ^160^ and ggtreeExtra ^161^ R packages.

Functional annotation of the MAGs was performed using the automated DRAM pipeline^162^. Additional databases used in the same DRAM run included the NCycDB ^163^, SCycDB ^164^, and PCycDB ^165^. KEGG annotations were integrated with annotations from NCycDB, SCycDB, and PCycDB based on KEGG Orthology (KO) numbers. This integration enabled differentiation between closely related orthologs from KEGG databases (e.g., K19044 encoding either amoA or pmoA) using metabolism-specific database results.

MAG stress tolerance trait identification was based on literature-derived definitions, KEGG database annotations ^166^, and modified microtrait definitions ^167^ as detailed in Supplementary Table 3. Traits were considered present in a MAG when at least 50% of trait- defining genes were detected. Functional pathway analysis for carbon, nitrogen, and sulfur metabolism in MAGs was conducted at both reaction (Supplementary Table 4) and pathway levels (Supplementary Table 5). Reaction and pathway definitions, including key enzymes, were sourced from the KEGG database and literature review. A reaction was considered present in a MAG when at least 50% of required genes were detected. Similarly, pathways were considered present when a MAG contained at least 50% of pathway reactions plus required key enzymes.

Maximal growth rate for each of the MAGs was predicted based on genome-wide codon usage statistics using the gRodon2 ^168^ R package.

### Metatranscriptomics processing

The quality of metatranscriptomics reads was evaluated with FastQC ^146^. Raw reads were processed with the “RQCFilter” pipeline (BBtools version 38.90) ^147^ to trim adapters and filter contaminants. Further filtering of risobomal RNA was performed using SortMeRNA (version 4.3.6) ^169^. Bowtie2 (v2.5.4) ^170^ was used to map the filtered and trimmed reads against the dereplicated MAGs. The resulting SAM files were sorted using samtools ^171^. Mapped reads counts were obtained using featureCounts (version 2.0.6) ^172^. Only transcripts with read counts higher than 5 and present in at least 3 samples were kept for downstream analysis. Read counts were normalized following the gene length corrected trimmed mean of M-values (geTMM) method ^173^.

### β-diversity analysis and ecological null modeling

#### Microbial phylogenetic signal

Microbial community assembly shifts during the monsoon season were analyzed using ecological null modeling ^43,44^,based on the assumption that phylogenetically related ASVs share ecological similarities (phylogenetic signal) ^44^. Phylogenetic signal analysis encompassed 19 environmental variables (detailed in Supplementary Table 1), using the between-ASV difference in environmental optima and between-ASV phylogenetic distance.

Environmental optima were calculated using abundance-weighted mean values for each environmental variable using the analogue package (version 0.17.6) ^174^. Between-ASV environmental optima differences were computed using Euclidean distance for all environmental variables. Between-OTU phylogenetic distances were derived from the microbial phylogenetic tree using the adephylo package (version 1.1.16) ^175^. Phylogenetic signal assessment was done using a Mantel correlogram based on a Pearson’s correlation coefficient across 50 distance classes with 999 permutations, applying progressive Holm-Bonferroni correction for multiple testing.

#### β-diversity analysis and ecological null modeling

Ecological processes influencing microbial and metabolome assemblages during monsoon periods were estimated using β-nearest taxon index (βNTI) and Raup-Crick Bray-Curtis index (RCBC), analyzing the microbial phylogenetic tree and TWCD metabolite-derived dendrogram as previously described ^43–45^.

Following this approach, |βNTI| > 2 indicates predominance of deterministic processes, such as environmental abiotic factors and biotic interactions influence microbial community turnover and differences in production and degradation rates of metabolites. Meanwhile, |βNTI| < 2 indicates that stochastic processes such as random processes of birth, death, colonization and speciation of microbial communities ^176^ cause changes in species diversity and composition. Similarly, metabolite shifts caused by physical forces or vector movements are considered as stochastic changes ^45^. Moreover, when βNTI > 2, variable selection indicates that divergent environmental factors drive high compositional turnover between a pair of communities. Conversely, when βNTI < -2, homogeneous selection indicates consistent selective pressures from stable environmental conditions are the primary cause of low compositional turnover between local communities ^43^.

Furthermore, the Raup-Crick Bray-Curtis (RCBC) turnover index was used to determine the influence of stochastic ecological processes. Briefly, the observed presence/absence-based Bray-Curtis values from pairwise community comparisons were calculated and compared to a null expectation (generated through 1,000 randomizations). Deviations of observed values from the null comparisons were then normalized between +1 and -1 to produce the RCBC metric. An RCBC > 0.95 with |βNTI| < 2 indicates that higher-than-expected compositional differences between communities (or metabolomes) are primarily due to dispersal limitation, allowing ecological drift. In contrast, an RCBC < -0.95 with |βNTI| < 2 suggests that lower-than-expected compositional differences are driven mainly by homogenizing dispersal. Finally, when |RCBC| < 0.95 and |βNTI| < 2, compositional turnover between communities (or metabolomes) is not predominantly influenced by selection, dispersal, or ecological drift—a condition referred to as ’undominated’ ^44^.

### Co-occurrence network analysis

MAG interactions were analyzed through co-occurrence microbial networks using a random matrix theory (RMT)-based approach ^70^. Networks were constructed using the Molecular Ecological Network Analysis pipeline (MENA) ^70,177^ website (http://ieg4.rccc.ou.edu/mena/login.cgi). Two separate networks were constructed, one using data from sampling points from two “dry” months (May and October) and another with data from two “wet” sampling points (July and August). Metagenome-assembled genomes abundances expressed as trimmed means of M-values (TMM) were obtained using CoverM. Separate abundance tables for each network were uploaded and processed using the MENA pipeline.

Briefly, a similarity matrix was built using Spearman correlations of features present in at least half of the samples. An association threshold was then determined using the RMT approach ^177^. Network topological parameters were calculated with the MENA pipeline. Module determination employed a greedy modularity optimization algorithm ^178^. Node topological roles were assigned based on within-module connectivity (Zi) and participation coefficient (Pi) ^70^. Networks were visualized using the igraph ^179^ and ggnetwork ^180^ R packages.

### Metabolic interaction network

Genome scale metabolic (GEM) models were reconstructed for each MAG using CarveMe ^181^, with distinct templates for bacterial and archaeal MAGs. Models were constructed without gap filling to avoid false-positive cross-feeding metabolic interaction predictions ^182^.

To better understand metabolic interactions among the microorganisms in each of the co-occurrence networks, metabolic interactions were assessed using SMETANA ^82^. SMETANA was run in “global” mode for each possible pair of MAGs within each of the two co-occurrence networks to calculate a metabolic interaction potential (MIP) score between each pair. Only pairs with MIP scores >= 5 were considered highly interacting and were used for building the metabolic interaction network ^183^. The cross feeding potential between pairs of highly interacting MAGS was then investigated using the “detailed” mode of SMETANA, keeping only those metabolites with a SMETANA score higher or equal than 0.1, as well as those related with nitrogen and sulfur metabolism. Node topological roles were assigned based on within-module connectivity (Zi) and participation coefficient (Pi) ^70^. Networks were visualized using the igraph and ggnetwork R packages.

## DATA AVAILABILITY

All environmental measurements are available as part of Supplementary Table 1.

Raw metagenomics, metatranscriptomics and amplicon sequencing data is available at the NCBI database (Bioproject Accession: PRJNA1193034). The following reviewer link can be used to check the submission until paper acceptance (https://dataview.ncbi.nlm.nih.gov/object/PRJNA1193034?reviewer=t661lr6tp95nc2n5fs8fi0fcai) Processing report of FTICR-MS data, assembled MAGs sequences and their annotations are available in the OSF repository with DOI (https://doi.org/10.17605/OSF.IO/WR6Z4)

## CODE AVAILABILITY

Code for all statistical analysis in this manuscript are available in its GitHub repository: https://github.com/Coayala/arid_dual_resilience_paper

## ACKNOWLEDGEMENTS

This study was supported in part by the University of Arizona (UA) UA Research, Innovation and Impact (RII) Core Facilities Pilot Program and UA Start-Up funds awarded to PI M. Tfaily. We thank the Environmental Molecular Sciences Laboratory (EMSL), a DOE Office of Science User Facility sponsored by the Office of Biological and Environmental Research and operated under Contract DE-AC05-76RL01830 and more specifically Rosalie Chu for their help in generating the FTICR-MS data. This includes EMSL project 60352 awarded to PI M. Tfaily. We thank the Arizona Genetics Core, University of Arizona, Tucson, AZ (<u>Facility RRID:SCR_012429</u>, https://azgc.arizona.edu/) for their help in generating all the next generation sequencing data.

We thank the Arizona Laboratory for Emerging Contaminants (ALEC), University of Arizona, Tucson, AZ, for their help with the analysis of total carbon, total nitrogen and metallic ions. Bioinformatics analysis in this publication used High Performance Computing (HPC) resources supported by the University of Arizona TRIF, UITS, and Research, Innovation, and Impact (RII) and maintained by the UArizona Research Technologies department. Additionally, this work used Bridges-2 at Pittsburgh Supercomputing Center through allocation BIO220147 from the Advanced Cyberinfrastructure Coordination Ecosystem: Services & Support (ACCESS) program, which is supported by U.S. National Science Foundation grants #2138259, #2138286, #2138307, #2137603, and #2138296.

## AUTHOR CONTRIBUTIONS

C.A.-O., V.F.-Z. and M.M.T conceptualized, designed and supervised the study. C.A.-O., V.F.-Z. and M.M.T. coordinated sampling efforts. C.A.-O. and V.F.-Z. collected samples. C.A.-O. and V.F.-Z. processed the samples. C.A.-O. and V.F.-Z. analyzed and visualized the data. C.A.-O., V.F.-Z. and M.M.T. drafted the paper. All authors provided comments and editions and approved of the final draft.

## ETHICS DECLARATIONS

### Competing Interest

The authors declare not competing interests

## Notes

https://doi.org/10.17605/OSF.IO/WR6Z4

https://dataview.ncbi.nlm.nih.gov/object/PRJNA1193034?reviewer=t661lr6tp95nc2n5fs8fi0fcai

### Competing Interest Statement

The authors have declared no competing interest.

## REFERENCES

1. Philippot, L., Chenu, C., Kappler, A., Rillig, M. C. & Fierer, N. The interplay between microbial communities and soil properties. Nat. Rev. Microbiol. 22, 226–239 (2024).

2. Nguyen, J., Lara-Gutiérrez, J. & Stocker, R. Environmental fluctuations and their effects on microbial communities, populations and individuals. FEMS Microbiol. Rev. 45, fuaa068 (2021).

3. Zhou, J. et al. Stochasticity, succession, and environmental perturbations in a fluidic ecosystem. Proc. Natl. Acad. Sci. U. S. A. 111, E836–45 (2014).

4. Philippot, L., Griffiths, B. S. & Langenheder, S. Microbial community resilience across ecosystems and multiple disturbances. Microbiol. Mol. Biol. Rev. 85, (2021).

5. Jansson, J. K. & Hofmockel, K. S. Soil microbiomes and climate change. Nat. Rev. Microbiol. 18, 35–46 (2020).

6. Shade, A. et al. Fundamentals of microbial community resistance and resilience. Front. Microbiol. 3, 417 (2012).

7. Allison, S. D. & Martiny, J. B. H. Colloquium paper: resistance, resilience, and redundancy in microbial communities. Proc. Natl. Acad. Sci. U. S. A. 105 **Suppl 1**, 11512–11519 (2008).

8. Schimel, J., Balser, T. C. & Wallenstein, M. Microbial stress-response physiology and its implications for ecosystem function. Ecology 88, 1386–1394 (2007).

9. Barnard, R. L., Osborne, C. & Firestone, M. Responses of soil bacterial and fungal communities to extreme desiccation and rewetting. ISME J. 7, 2229–2241 (2013).

10. Schimel, J. P. Life in dry soils: Effects of drought on soil microbial communities and processes. Annu. Rev. Ecol. Evol. Syst. 49, 409–432 (2018).

11. Ouyang, Y. & Li, X. Recent research progress on soil microbial responses to drying– rewetting cycles. Sheng Tai Xue Bao 33, 1–6 (2013).

12. Bardgett, R. D. & Caruso, T. Soil microbial community responses to climate extremes: resistance, resilience and transitions to alternative states. Philos. Trans. R. Soc. Lond. B Biol. Sci. 375, 20190112 (2020).

13. Collins, S. L. et al. A multiscale, hierarchical model of pulse dynamics in arid-land ecosystems. Annu. Rev. Ecol. Evol. Syst. 45, 397–419 (2014).

14. Maestre, F. T. et al. Structure and functioning of dryland ecosystems in a changing world. Annu. Rev. Ecol. Evol. Syst. 47, 215–237 (2016).

15. Jiang, H. et al. Seasonal dynamics of soil microbiome in response to dry-wet alternation along the Jinsha River Dry-hot Valley. BMC Microbiol. 24, 496 (2024).

16. Naidoo, Y., Valverde, A., Pierneef, R. E. & Cowan, D. A. Differences in precipitation regime shape microbial community composition and functional potential in Namib Desert soils. Microb. Ecol. 83, 689–701 (2022).

17. Overpeck, J. T. & Udall, B. Climate change and the aridification of North America. Proc. Natl. Acad. Sci. U. S. A. 117, 11856–11858 (2020).

18. Burrell, A. L., Evans, J. P. & De Kauwe, M. G. Anthropogenic climate change has driven over 5 million km2 of drylands towards desertification. Nat. Commun. 11, 3853 (2020).

19. Meier, D. V., Imminger, S., Gillor, O. & Woebken, D. Distribution of mixotrophy and desiccation survival mechanisms across microbial genomes in an arid biological soil crust community. mSystems 6, (2021).

20. Imminger, S. et al. Survival and rapid resuscitation permit limited productivity in desert microbial communities. Nat. Commun. 15, 1–17 (2024).

21. Shoemaker, W. R. & Lennon, J. T. Evolution with a seed bank: The population genetic consequences of microbial dormancy. Evol. Appl. 11, 60–75 (2018).

22. León-Sobrino, C., Ramond, J.-B., Maggs-Kölling, G. & Cowan, D. A. Nutrient Acquisition, Rather Than Stress Response Over Diel Cycles, Drives Microbial Transcription in a Hyper-Arid Namib Desert Soil. Front. Microbiol. 10, 1054 (2019).

23. Coclet, C., Cowan, D. & Lebre, P. H. Survival under Stress: Microbial Adaptation in Hot Desert Soils. in Microbiology of Hot Deserts (eds. Ramond, J.-B. & Cowan, D. A.) 293–317 (Springer International Publishing, Cham, 2022).

24. Maestre, F. T. et al. Research needs on the biodiversity-ecosystem functioning relationship in drylands. NPJ Biodivers 3, 12 (2024).

25. Zolotokrylin, A. N., Titkova, T. B. & Brito-Castillo, L. Wet and dry patterns associated with ENSO events in the Sonoran Desert from, 2000–2015. J. Arid Environ. 134, 21–32 (2016).

26. Adams, D. K. & Comrie, A. C. The North American Monsoon. Bull. Am. Meteorol. Soc. 78, 2197–2214 (1997).

27. Li, C. et al. The adjustment of life history strategies drives the ecological adaptations of soil microbiota to aridity. Mol. Ecol. 31, 2920–2934 (2022).

28. Evans, S. E. & Wallenstein, M. D. Soil microbial community response to drying and rewetting stress: does historical precipitation regime matter? Biogeochemistry 109, 101– 116 (2012).

29. Wallenstein, M. D. & Hall, E. K. A trait-based framework for predicting when and where microbial adaptation to climate change will affect ecosystem functioning. Biogeochemistry 109, 35–47 (2012).

30. Wu, K., Xu, W. & Yang, W. Effects of precipitation changes on soil bacterial community composition and diversity in the Junggar desert of Xinjiang, China. PeerJ 8, e8433 (2020).

31. Vásquez-Dean, J., Maza, F., Morel, I., Pulgar, R. & González, M. Microbial communities from arid environments on a global scale. A systematic review. Biol. Res. 53, 29 (2020).

32. Šťovíček, A., Kim, M., Or, D. & Gillor, O. Microbial community response to hydration-desiccation cycles in desert soil. Sci. Rep. 7, 45735 (2017).

33. Blazewicz, S. J., Schwartz, E. & Firestone, M. K. Growth and death of bacteria and fungi underlie rainfall-induced carbon dioxide pulses from seasonally dried soil. Ecology 95, 1162–1172 (2014).

34. Lindström, E. S. & Langenheder, S. Local and regional factors influencing bacterial community assembly: Bacterial community assembly. Environ. Microbiol. Rep. 4, 1–9 (2012).

35. Nemergut, D. R. et al. Patterns and processes of microbial community assembly. Microbiol. Mol. Biol. Rev. 77, 342–356 (2013).

36. Wilson, R. M. et al. Soil metabolome response to whole-ecosystem warming at the Spruce and Peatland Responses under Changing Environments experiment. Proc. Natl. Acad. Sci. U. S. A. 118, (2021).

37. AminiTabrizi, R. et al. Controls on Soil Organic Matter Degradation and Subsequent Greenhouse Gas Emissions Across a Permafrost Thaw Gradient in Northern Sweden. Front Earth Sci. Chin. 8, (2020).

38. Ramond, J.-B. & Cowan, D. A. Microbial ecology of hot desert soils. in Ecological Studies 89–110 (Springer International Publishing, Cham, 2022).

39. Moyano, F. E. et al. The moisture response of soil heterotrophic respiration: interaction with soil properties. Biogeosciences 9, 1173–1182 (2012).

40. Yu, Z. et al. Molecular insights into the transformation of dissolved organic matter during hyperthermophilic composting using ESI FT-ICR MS. Bioresour. Technol. 292, 122007 (2019).

41. Li, H. et al. Responses of soil bacterial communities to nitrogen deposition and precipitation increment are closely linked with aboveground community variation. Microb. Ecol. 71, 974– 989 (2016).

42. Chen, Q., Niu, B., Hu, Y., Luo, T. & Zhang, G. Warming and increased precipitation indirectly affect the composition and turnover of labile-fraction soil organic matter by directly affecting vegetation and microorganisms. Sci. Total Environ. 714, 136787 (2020).

43. Stegen, J. C., Lin, X., Fredrickson, J. K. & Konopka, A. E. Estimating and mapping ecological processes influencing microbial community assembly. Front. Microbiol. 6, 370 (2015).

44. Stegen, J. C., Lin, X., Konopka, A. E. & Fredrickson, J. K. Stochastic and deterministic assembly processes in subsurface microbial communities. ISME J. 6, 1653–1664 (2012).

45. Danczak, R. E. et al. Using metacommunity ecology to understand environmental metabolomes. Nat. Commun. 11, 6369 (2020).

46. Freire-Zapata, V. et al. Microbiome-metabolite linkages drive greenhouse gas dynamics over a permafrost thaw gradient. Nat. Microbiol. 1–17 (2024).

47. 47. Wang, X.-B., et al. A drying-rewetting cycle imposes more important shifts on soil microbial communities than does reduced precipitation. mSystems 7, e0024722 (2022).

48. Zhou, J. & Ning, D. Stochastic community assembly: Does it matter in microbial ecology? Microbiol. Mol. Biol. Rev. 81, (2017).

49. Plaza, C., Gascó, G., Méndez, A. M., Zaccone, C. & Maestre, F. T. Soil organic matter in dryland ecosystems. in The Future of Soil Carbon 39–70 (Elsevier, 2018).

50. Kim, M. & Or, D. Individual-based model of microbial life on hydrated rough soil surfaces. PLoS One 11, e0147394 (2016).

51. Warren, C. R. Response of osmolytes in soil to drying and rewetting. Soil Biol. Biochem. 70, 22–32 (2014).

52. Halverson, L. J., Jones, T. M. & Firestone, M. K. Release of intracellular solutes by four soil bacteria exposed to dilution stress. Soil Sci. Soc. Am. J. 64, 1630–1637 (2000).

53. Martínez-Mena, M., Alvarez Rogel, J., Castillo, V. & Albaladejo, J. Organic carbon and nitrogen losses influenced by vegetation removal in a semiarid mediterranean soil. Biogeochemistry 61, 309–321 (2002).

54. Taravati, A. et al. Various effects of sugar and polyols on the protein structure and function: role as osmolyte on protein stability. World Appl Sci J 2, 353–362 (2007).

55. Couvillion, S. P. et al. Rapid remodeling of the soil lipidome in response to a drying- rewetting event. Microbiome 11, 34 (2023).

56. Lal, R. Carbon sequestration in dryland ecosystems. Environ. Manage. 33, 528–544 (2004).

57. Liu, L. et al. Microbial diversity and adaptive strategies in the Mars-like Qaidam Basin, North Tibetan Plateau, China. Environ. Microbiol. Rep. 14, 873–885 (2022).

58. Musto, H. et al. Correlations between genomic GC levels and optimal growth temperatures in prokaryotes. FEBS Lett. 573, 73–77 (2004).

59. Fierer, N. et al. Cross-biome metagenomic analyses of soil microbial communities and their functional attributes. Proc. Natl. Acad. Sci. U. S. A. 109, 21390–21395 (2012).

60. Chuckran, P. F. et al. Edaphic controls on genome size and GC content of bacteria in soil microbial communities. Soil Biol. Biochem. 178, 108935 (2023).

61. Giovannoni, S. J., Cameron Thrash, J. & Temperton, B. Implications of streamlining theory for microbial ecology. ISME J. 8, 1553–1565 (2014).

62. Liu, Q., Chen, Y. & Xu, X.-W. Genomic insight into strategy, interaction and evolution of nitrifiers in metabolizing key labile-dissolved organic nitrogen in different environmental niches. Front. Microbiol. 14, 1273211 (2023).

63. Ranawat, P. & Rawat, S. Stress response physiology of thermophiles. Arch. Microbiol. 199, 391–414 (2017).

64. Sun, D. et al. KatG and KatE confer Acinetobacter resistance to hydrogen peroxide but sensitize bacteria to killing by phagocytic respiratory burst. Life Sci. 148, 31–40 (2016).

65. Karas, V. O., Westerlaken, I. & Meyer, A. S. The DNA-binding protein from starved cells (DPs) utilizes dual functions to defend cells against multiple stresses. J. Bacteriol. 197, 3206–3215 (2015).

66. Ahn, S., Jung, J., Jang, I.-A., Madsen, E. L. & Park, W. Role of glyoxylate shunt in oxidative stress response. J. Biol. Chem. 291, 11928–11938 (2016).

67. Kuzminov, A. Recombinational Repair of DNA Damage in *Escherichia coli* and Bacteriophage λ. Microbiol. Mol. Biol. Rev. 63, 751–813 (1999).

68. Zahradka, K. et al. Reassembly of shattered chromosomes in Deinococcus radiodurans. Nature 443, 569–573 (2006).

69. Leem, S. H., Ropp, P. A. & Sugino, A. The yeast Saccharomyces cerevisiae DNA polymerase IV: possible involvement in double strand break DNA repair. Nucleic Acids Res. 22, 3011–3017 (1994).

70. Deng, Y. et al. Molecular ecological network analyses. BMC Bioinformatics 13, 113 (2012).

71. 71. Wang, X., et al. Decreased soil multifunctionality is associated with altered microbial network properties under precipitation reduction in a semiarid grassland. Imeta 2, e106 (2023).

72. Wang, S., Wang, X., Han, X. & Deng, Y. Higher precipitation strengthens the microbial interactions in semi-arid grassland soils. Glob. Ecol. Biogeogr. 27, 570–580 (2018).

73. Evans, S. E. & Wallenstein, M. D. Climate change alters ecological strategies of soil bacteria. Ecol. Lett. 17, 155–164 (2014).

74. 74. Leung, P. M., et al. Energetic basis of microbial growth and persistence in desert ecosystems. mSystems 5, (2020).

75. Hu, Y. et al. Seasonal patterns of soil microbial community response to warming and increased precipitation in a semiarid steppe. Appl. Soil Ecol. 182, 104712 (2023).

76. Liu, D. et al. Response of microbial communities and their metabolic functions to Drying^−^Rewetting stress in a temperate forest soil. Microorganisms 7, 129 (2019).

77. Nielsen, U. N. & Ball, B. A. Impacts of altered precipitation regimes on soil communities and biogeochemistry in arid and semi-arid ecosystems. Glob. Chang. Biol. 21, 1407–1421 (2015).

78. Armstrong, A. et al. Temporal dynamics of hot desert microbial communities reveal structural and functional responses to water input. Sci. Rep. 6, 34434 (2016).

79. Marasco, R. et al. Rhizosheath microbial community assembly of sympatric desert speargrasses is independent of the plant host. Microbiome 6, 215 (2018).

80. Mandakovic, D. et al. Structure and co-occurrence patterns in microbial communities under acute environmental stress reveal ecological factors fostering resilience. Sci. Rep. 8, 5875 (2018).

81. Mosqueira, M. J. et al. Consistent bacterial selection by date palm root system across heterogeneous desert oasis agroecosystems. Sci. Rep. 9, 4033 (2019).

82. Zelezniak, A. et al. Metabolic dependencies drive species co-occurrence in diverse microbial communities. Proc. Natl. Acad. Sci. U. S. A. 112, 6449–6454 (2015).

83. Kpalari, D. F. et al. Soil bacterial community and greenhouse gas emissions as responded to the coupled application of nitrogen fertilizer and microbial decomposing inoculants in wheat (Triticum aestivum L.) seedling stage under different water regimes. Agronomy (Basel) 13, 2950 (2023).

84. Yang, Q. et al. Erosion and deposition significantly affect the microbial diversity, co- occurrence network, and multifunctionality in agricultural soils of Northeast China. J. Soils Sediments 24, 888–900 (2023).

85. He, J. et al. Distinct composition patterns of bacterial and fungal communities and biogeochemical cycling genes depend on the vegetation type in arid soil. Appl. Soil Ecol. 191, 105064 (2023).

86. Li, J., Liu, T., McIlroy, S. J., Tyson, G. W. & Guo, J. Phylogenetic and metabolic diversity of microbial communities performing anaerobic ammonium and methane oxidations under different nitrogen loadings. ISME Commun. 3, 39 (2023).

87. Mee, M. T., Collins, J. J., Church, G. M. & Wang, H. H. Syntrophic exchange in synthetic microbial communities. Proc. Natl. Acad. Sci. U. S. A. 111, E2149–56 (2014).

88. Shen, J.-P., Xu, Z. & He, J.-Z. Frontiers in the microbial processes of ammonia oxidation in soils and sediments. J. Soils Sediments 14, 1023–1029 (2014).

89. Ramond, J.-B., Jordaan, K., Díez, B., Heinzelmann, S. M. & Cowan, D. A. Microbial Biogeochemical Cycling of Nitrogen in Arid Ecosystems. Microbiol. Mol. Biol. Rev. 86, e00109–21 (2022).

90. Bei, Q. et al. Metabolic potential of Nitrososphaera-associated clades. ISME J. 18, wrae086 (2024).

91. León-Sobrino, C. et al. Temporal dynamics of microbial transcription in wetted hyperarid desert soils. FEMS Microbiol. Ecol. 100, (2024).

92. Lu, L. & Jia, Z. Urease gene-containing Archaea dominate autotrophic ammonia oxidation in two acid soils: Urea-linked archaeal ammonia oxidation in acid soil. Environ. Microbiol. 15, 1795–1809 (2013).

93. Hatzenpichler, R. Diversity, physiology, and niche differentiation of ammonia-oxidizing archaea. Appl. Environ. Microbiol. 78, 7501–7510 (2012).

94. Zhao, R., Wang, H., Yang, H., Yun, Y. & Barton, H. A. Ammonia-oxidizing Archaea dominate ammonia-oxidizing communities within alkaline cave sediments. Geomicrobiol. J. 34, 511–523 (2017).

95. Gao, S. et al. Effects of temperature increase and nitrogen addition on the early litter decomposition in permafrost peatlands. Catena 209, 105801 (2022).

96. Neilson, J. W. et al. Significant impacts of increasing aridity on the arid soil microbiome. mSystems 2, (2017).

97. Yuan, M. M. et al. Climate warming enhances microbial network complexity and stability. Nat. Clim. Chang. 11, 343–348 (2021).

98. Bertness, M. D. & Callaway, R. Positive interactions in communities. Trends in ecology & evolution 9, 191–193 (1994).

99. Hammarlund, S. P. & Harcombe, W. R. Refining the stress gradient hypothesis in a microbial community. Proc. Natl. Acad. Sci. U. S. A. 116, 15760–15762 (2019).

100. Hernandez, D. J., David, A. S., Menges, E. S., Searcy, C. A. & Afkhami, M. E. Environmental stress destabilizes microbial networks. ISME J. 15, 1722–1734 (2021).

101. de Vries, F. T. et al. Soil bacterial networks are less stable under drought than fungal networks. Nat. Commun. 9, 3033 (2018).

102. Goberna, M. et al. Incorporating phylogenetic metrics to microbial co-occurrence networks based on amplicon sequences to discern community assembly processes. Mol. Ecol. Resour. 19, 1552–1564 (2019).

103. Jia, M. et al. Effects of precipitation change and nitrogen addition on the composition, diversity, and molecular ecological network of soil bacterial communities in a desert steppe. PLoS One 16, e0248194 (2021).

104. Gao, C. et al. Co-occurrence networks reveal more complexity than community composition in resistance and resilience of microbial communities. Nat. Commun. 13, 3867 (2022).

105. Zomorrodi, A. R. & Segrè, D. Genome-driven evolutionary game theory helps understand the rise of metabolic interdependencies in microbial communities. Nat. Commun. 8, 1563 (2017).

106. Tfaily, M. M., Hess, N. J., Koyama, A. & Evans, R. D. Elevated [CO2] changes soil organic matter composition and substrate diversity in an arid ecosystem. Geoderma 330, 1–8 (2018).

107. Honeker, L. K. et al. Elucidating drought-tolerance mechanisms in plant roots through 1H NMR metabolomics in parallel with MALDI-MS, and NanoSIMS imaging techniques. Environ. Sci. Technol. 56, 2021–2032 (2022).

108. Shivlata, L. & Satyanarayana, T. Thermophilic and alkaliphilic Actinobacteria: biology and potential applications. Front. Microbiol. 6, 1014 (2015).

109. Hamedi, J., Poorinmohammad, N. & Papiran, R. Growth and Life Cycle of Actinobacteria. in Biology and Biotechnology of Actinobacteria (eds. Wink, J., Mohammadipanah, F. & Hamedi, J.) 29–50 (Springer International Publishing, Cham, 2017).

110. Bouskill, N. J. et al. Belowground response to drought in a tropical forest soil. I. changes in microbial functional potential and metabolism. Front. Microbiol. 7, 525 (2016).

111. Ramin, K. I. & Allison, S. D. Bacterial tradeoffs in growth rate and extracellular enzymes. Front. Microbiol. 10, 2956 (2019).

112. Pointing, S. B. & Belnap, J. Microbial colonization and controls in dryland systems. Nat. Rev. Microbiol. 10, 551–562 (2012).

113. Ochoa-Hueso, R. et al. Drought consistently alters the composition of soil fungal and bacterial communities in grasslands from two continents. Glob. Chang. Biol. 24, 2818–2827 (2018).

114. Veach, A. M. & Zeglin, L. H. Historical drought affects microbial population dynamics and activity during soil drying and re-wet. Microb. Ecol. 79, 662–674 (2020).

115. Ouyang, Y. & Li, X. Effect of repeated drying-rewetting cycles on soil extracellular enzyme activities and microbial community composition in arid and semi-arid ecosystems. Eur. J. Soil Biol. 98, 103187 (2020).

116. Weiss, J. L. & Overpeck, J. T. Is the Sonoran Desert losing its cool? Glob. Chang. Biol. 11, 2065–2077 (2005).

117. Biederman, J. A. et al. Shrubland carbon sink depends upon winter water availability in the warm deserts of North America. Agric. For. Meteorol. 249, 407–419 (2018).

118. Douglas, M., Maddox, R., Howard, K. & Reyes, S. The Mexican monsoon. Journal of Climate 6, 1665–1677 (1993).

119. AminiTabrizi, R., Dontsova, K., Graf Grachet, N. & Tfaily, M. M. Elevated temperatures drive abiotic and biotic degradation of organic matter in a peat bog under oxic conditions. Sci. Total Environ. 804, 150045 (2022).

120. 120. Didan, K. MODIS/Terra Vegetation Indices 16-Day L3 Global 250m SIN Grid V061 [Data set]. 10.5067/MODIS/MOD13Q1.061 (2021).

121. Dittmar, T., Koch, B., Hertkorn, N. & Kattner, G. A simple and efficient method for the solid- phase extraction of dissolved organic matter (SPE-DOM) from seawater: SPE-DOM from seawater. Limnol. Oceanogr. Methods 6, 230–235 (2008).

122. Tolić, N. et al. Formularity: Software for automated formula assignment of natural and other organic matter from ultrahigh-resolution mass spectra. Anal. Chem. 89, 12659–12665 (2017).

123. Ayala-Ortiz, C. et al. MetaboDirect: an analytical pipeline for the processing of FT-ICR MS- based metabolomic data. Microbiome 11, 28 (2023).

124. Tfaily, M. M. et al. Sequential extraction protocol for organic matter from soils and sediments using high resolution mass spectrometry. Anal. Chim. Acta 972, 54–61 (2017).

125. Klindworth, A. et al. Evaluation of general 16S ribosomal RNA gene PCR primers for classical and next-generation sequencing-based diversity studies. Nucleic Acids Res. 41, e1 (2013).

126. Bay, S. K. et al. Chemosynthetic and photosynthetic bacteria contribute differentially to primary production across a steep desert aridity gradient. ISME J. 15, 3339–3356 (2021).

127. Lian, W.-H. et al. Culturomics- and metagenomics-based insights into the microbial community and function of rhizosphere soils in Sinai desert farming systems. *Environ*. Microbiome 18, 4 (2023).

128. Zhao, M. et al. Asymmetric responses of abundance and diversity of N-cycling genes to altered precipitation in arid grasslands. Funct. Ecol. 37, 2953–2966 (2023).

129. Nguyen, T. M. et al. Whole community shotgun metagenomes of two biological soil crust types from the Mojave Desert. Microbiol. Resour. Announc. 13, e0098023 (2024).

130. Barrón-Sandoval, A. et al. Functional significance of microbial diversity in arid soils: biological soil crusts and nitrogen fixation as a model system. FEMS Microbiol. Ecol. 99, fiad009 (2023).

131. Chen, Y., Neilson, J. W., Kushwaha, P., Maier, R. M. & Barberán, A. Life-history strategies of soil microbial communities in an arid ecosystem. ISME J. 15, 649–657 (2021).

132. Ondov, B. D. et al. Mash: fast genome and metagenome distance estimation using MinHash. Genome Biol. 17, 132 (2016).

133. Woodcroft, B. J., et al. SingleM and Sandpiper: Robust microbial taxonomic profiles from metagenomic data. *bioRxiv* 2024.01.30.578060 (2024) doi:10.1101/2024.01.30.578060.

134. Callahan, B. J. et al. DADA2: High-resolution sample inference from Illumina amplicon data. Nat. Methods 13, 581–583 (2016).

135. Martin, M. Cutadapt removes adapter sequences from high-throughput sequencing reads. EMBnet J. 17, 10 (2011).

136. Wang, Q., Garrity, G. M., Tiedje, J. M. & Cole, J. R. Naïve Bayesian classifier for rapid assignment of rRNA sequences into the new bacterial taxonomy. Appl. Environ. Microbiol. 73, 5261–5267 (2007).

137. Parks, D. H. et al. GTDB: an ongoing census of bacterial and archaeal diversity through a phylogenetically consistent, rank normalized and complete genome-based taxonomy.Nucleic Acids Res. 50, D785–D794 (2022).

138. Větrovský, T. & Baldrian, P. The variability of the 16S rRNA gene in bacterial genomes and its consequences for bacterial community analyses. PLoS One 8, e57923 (2013).

139. 139. Lan, Y., Rosen, G. & Hershberg, R. Marker genes that are less conserved in their sequences are useful for predicting genome-wide similarity levels between closely related prokaryotic strains. Microbiome 4, 18 (2016).

140. 140. R Core Team. R: A Language and Environment for Statistical Computing. Preprint at https://www.R-project.org/ (2023).

141. McMurdie, P. J. & Holmes, S. phyloseq: an R package for reproducible interactive analysis and graphics of microbiome census data. PLoS One 8, e61217 (2013).

142. 142. Oksanen, J., et al. Vegan: Community Ecology Package. https://github.com/vegandevs/vegan (2024).

143. Wright, E. Using DECIPHER v2.0 to analyze big biological sequence data in R. R J. 8, 352 (2016).

144. Price, M. N., Dehal, P. S. & Arkin, A. P. FastTree 2--approximately maximum-likelihood trees for large alignments. PLoS One 5, e9490 (2010).

145. Revell, L. J. phytools 2.0: an updated R ecosystem for phylogenetic comparative methods (and other things). PeerJ 12, e16505 (2024).

146. 146. Andrews, S. Babraham Bioinformatics - FastQC A Quality Control tool for High Throughput Sequence Data. FastQC https://www.bioinformatics.babraham.ac.uk/projects/fastqc/ (2010).

147. 147. Bushnell, B. BBMap. SourceForge https://sourceforge.net/projects/bbmap/ (2021).

148. 148. 148.Li, D., et al. MEGAHIT v1.0: A fast and scalable metagenome assembler driven by advanced methodologies and community practices. Methods 102, 3–11 (2016).

149. Aroney, S. T. N., et al. CoverM: Read Coverage Calculator for Metagenomics. (2024). doi:10.5281/zenodo.10531253.

150. Alneberg, J. et al. Binning metagenomic contigs by coverage and composition. Nat. Methods 11, 1144–1146 (2014).

151. Kang, D. D. et al. MetaBAT 2: an adaptive binning algorithm for robust and efficient genome reconstruction from metagenome assemblies. PeerJ 7, e7359 (2019).

152. Wu, Y.-W., Simmons, B. A. & Singer, S. W. MaxBin 2.0: an automated binning algorithm to recover genomes from multiple metagenomic datasets. Bioinformatics 32, 605–607 (2016).

153. Uritskiy, G. V., DiRuggiero, J. & Taylor, J. MetaWRAP-a flexible pipeline for genome- resolved metagenomic data analysis. Microbiome 6, 158 (2018).

154. Eren, A. M. et al. Anvi’o: an advanced analysis and visualization platform for ’omics data. PeerJ 3, e1319 (2015).

155. Chklovski, A., Parks, D. H., Woodcroft, B. J. & Tyson, G. W. CheckM2: a rapid, scalable and accurate tool for assessing microbial genome quality using machine learning. Nat. Methods 20, 1203–1212 (2023).

156. Jain, C., Rodriguez-R, L. M., Phillippy, A. M., Konstantinidis, K. T. & Aluru, S. High throughput ANI analysis of 90K prokaryotic genomes reveals clear species boundaries. Nat. Commun. 9, 5114 (2018).

157. Woodcroft, B. J. Galah: More Scalable Dereplication for Metagenome Assembled Genomes. (Github, 2024).

158. Xu, L. et al. Genome-resolved metagenomics reveals role of iron metabolism in drought- induced rhizosphere microbiome dynamics. Nat. Commun. 12, 3209 (2021).

159. Chaumeil, P.-A., Mussig, A. J., Hugenholtz, P. & Parks, D. H. GTDB-Tk v2: memory friendly classification with the genome taxonomy database. Bioinformatics 38, 5315–5316 (2022).

160. Yu, G., Smith, D. K., Zhu, H., Guan, Y. & Lam, T. T.-Y. Ggtree: An r package for visualization and annotation of phylogenetic trees with their covariates and other associated data. Methods Ecol. Evol. 8, 28–36 (2017).

161. Xu, S. et al. GgtreeExtra: Compact visualization of richly annotated phylogenetic data. Mol. Biol. Evol. 38, 4039–4042 (2021).

162. Shaffer, M. et al. DRAM for distilling microbial metabolism to automate the curation of microbiome function. Nucleic Acids Res. 48, 8883–8900 (2020).

163. Tu, Q., Lin, L., Cheng, L., Deng, Y. & He, Z. NCycDB: a curated integrative database for fast and accurate metagenomic profiling of nitrogen cycling genes. Bioinformatics 35, 1040–1048 (2019).

164. Yu, X. et al. SCycDB: A curated functional gene database for metagenomic profiling of sulphur cycling pathways. Mol. Ecol. Resour. 21, 924–940 (2021).

165. Zeng, J. et al. PCycDB: a comprehensive and accurate database for fast analysis of phosphorus cycling genes. Microbiome 10, 101 (2022).

166. Kanehisa, M. The KEGG database. *Novartis Found. Symp*. 247, 91–101; discussion 101–3, 119–28, 244–52 (2002).

167. Karaoz, U. & Brodie, E. L. MicroTrait: A toolset for a trait-based representation of microbial genomes. Front. Bioinform. 2, 918853 (2022).

168. Weissman, J. L., Hou, S. & Fuhrman, J. A. Estimating maximal microbial growth rates from cultures, metagenomes, and single cells via codon usage patterns. Proc. Natl. Acad. Sci. U. S. A. 118, e2016810118 (2021).

169. Kopylova, E., Noé, L. & Touzet, H. SortMeRNA: fast and accurate filtering of ribosomal RNAs in metatranscriptomic data. Bioinformatics 28, 3211–3217 (2012).

170. Langmead, B. & Salzberg, S. L. Fast gapped-read alignment with Bowtie 2. Nat. Methods 9, 357–359 (2012).

171. Li, H. et al. The Sequence Alignment/Map format and SAMtools. Bioinformatics 25, 2078– 2079 (2009).

172. Liao, Y., Smyth, G. K. & Shi, W. featureCounts: an efficient general purpose program for assigning sequence reads to genomic features. Bioinformatics 30, 923–930 (2014).

173. Smid, M. et al. Gene length corrected trimmed mean of M-values (GeTMM) processing of RNA-seq data performs similarly in intersample analyses while improving intrasample comparisons. BMC Bioinformatics 19, 236 (2018).

174. Simpson, G. L. Analogue Methods in Palaeoecology: Using theanaloguePackage. J. Stat. Softw. 22, 1–29 (2007).

175. Jombart, T., Balloux, F. & Dray, S. Adephylo: New tools for investigating the phylogenetic signal in biological traits. Bioinformatics 26, 1907–1909 (2010).

176. Chase, J. M. Drought mediates the importance of stochastic community assembly. Proc. Natl. Acad. Sci. U. S. A. 104, 17430–17434 (2007).

177. Zhou, J. et al. Functional molecular ecological networks. MBio 1, (2010).

178. Newman, M. E. J. Modularity and community structure in networks. Proc. Natl. Acad. Sci. U. S. A. 103, 8577–8582 (2006).

179. Csardi, G. & Nepusz, T. The igraph software package for complex network research. Complex Systems 1695 (2005).

180. 180. Briatte, F. ggnetwork: Geometries to Plot Networks with ‘ggplot2’. Preprint at https://CRAN.R-project.org/package=ggnetwork (2024).

181. Machado, D., Andrejev, S., Tramontano, M. & Patil, K. R. Fast automated reconstruction of genome-scale metabolic models for microbial species and communities. Nucleic Acids Res. 46, 7542–7553 (2018).

182. Giordano, N. et al. Genome-scale community modelling reveals conserved metabolic cross- feedings in epipelagic bacterioplankton communities. Nat. Commun. 15, 2721 (2024).

183. Du, H. et al. Microbial active functional modules derived from network analysis and metabolic interactions decipher the complex microbiome assembly in mangrove sediments. Microbiome 10, 224 (2022).

