## Supplementary Note for "Stochastic Assembly and Metabolic Network Reorganization Drive Microbial Resilience in Arid Soils"

### SUPPLEMENTARY NOTES

#### Supplementary Note 1

#### Community Structure and Metabolite Dynamics During Monsoon Transitions in Arid Soils

*This note characterizes the taxonomic composition and metabolite profiles of arid soil microbiomes throughout a monsoon cycle, integrating both amplicon sequencing and shotgun metagenomic approaches with high-resolution metabolomics. The analysis reveals how dominant taxa (Actinomycetota, Pseudomonadota, and Thermoproteota) employ diverse stress tolerance mechanisms, while metabolite profiles show distinct temporal patterns reflecting both microbial mortality and adaptation during wet-dry transitions. The findings demonstrate how community structure and metabolic responses are coordinated across multiple organizational levels to maintain ecosystem function under extreme environmental fluctuations.*

##### Taxonomic profile

Taxonomic analysis revealed a diverse, yet specialized community adapted to arid conditions. We identified 6,698 unique amplicon sequencing variants (ASVs) through 16S rRNA sequencing and 807 operational taxonomic units (OTUs) via SingleM analysis [1] of shotgun metagenomics data. The community was dominated by Actinomycetota (ASV: 50.1%; SingleM: 64.5%) and Pseudomonadota (ASV: 17.6%; SingleM: 8.9%), followed by Acidobacteriota and Chloroflexota (Fig 2A and Supplementary Fig 2A), consistent with previous observations in southwestern USA [2,3], and other arid systems [4–9]. The prevalence of these taxa reflects diverse individual-level adaptations that support community resilience.Actinomycetota's dominance stems from multiple stress tolerance mechanisms: extreme temperature and UV radiation resistance, metabolic versatility, sporulation capabilities, and efficient DNA repair strategies [10–13]. Pseudomonadota (previously Proteobacteria) persist through bacteriochlorophyll-dependent photosynthesis, enabling survival in nutrient-poor conditions [6]. Chloroflexota's success appears linked to their slow growth rates [14], an adaptation to resource limitation and drought stress [9,15].

Interestingly, Thermoproteota showed marked differences between detection methods, representing 3.5% of SingleM OTUs but <0.1% of ASVs. Thermoproteota (previously known as Thaumarchaeota), is an archaeal taxon previously documented in various hot desert soils such as the Mojave and the Chihuahuan Desert [16], that plays a key role in arid [17–20] and oligotrophic environments [21] through ammonia oxidation and carbon fixation [17]. The detection discrepancy between methods likely reflects methodological biases including 16S copy number variation,

PCR amplification biases, and differential primer specificity [22], highlighting the importance of complementary approaches for comprehensive community characterization.

These findings reinforce our dual-level resilience framework by demonstrating how diverse individual adaptations collectively maintain community stability in extreme conditions. The consistent community structure across geographically distinct arid environments suggests fundamental constraints on microbial adaptation strategies, while methodological insights inform future studies of arid ecosystem resilience.

### Organic matter profile

Fourier transform ion cyclotron resonance mass spectrometry (FTICR-MS) analysis in positive mode revealed a complex metabolic landscape reflecting both community-level processes and individual adaptations. We identified 20,971 distinct masses (m/z) across all samples, with 4,175 assigned putative molecular formulas using Formularity [23]. The metabolite profile was dominated by lipid-like compounds, followed by protein- and lignin-like metabolites (Supplementary Fig. 2d), contrasting with peatlands [24–26] and forest ecosystems [27],[28], [29] where plant-derived lignin dominates [30]. This distinct profile reflects the primacy of microbial rather than plant influences on biogeochemical processes in arid soils [31], [32].

Elemental composition analysis revealed a high abundance of N-containing (CHON, CHONS, CHONP, CHONSP) and S-containing compounds (CHOS, CHONS, CHOSP, CHONSP). The prevalence of N-containing metabolites indicates substantial microbial contribution to soil organic matter (SOM) formation [33]through protein degradation and biomass turnover [34]. S-containing compounds likely derive from multiple sources: atmospheric sulfate deposition [35], urban particulate matter [36], Cenozoic evaporites [37] and anthropogenic inputs [29,38,39]. The widespread distribution of assimilatory sulfate reduction pathways among MAGs (Supplementary Note 2, Supplementary Fig. 4) demonstrates active microbial incorporation of these compounds.

While the abundance of some lipid-like compounds remained high throughout the monsoon season (Supplementary Fig. 2h), other metabolites such as some amino sugars increased in abundance alongside the increase in water availability. Amino sugars are highly labile metabolites [40], derived from microbial components [33], that can be used as a marker of microbial necromass [41]. The increase of some amino sugars, especially at the beginning of the monsoon season, is a potential indicator of how the sudden increase of water may result in the dead of some microbial communities due to osmotic pressure or oxygen limitation [42,43], increasing the nutrient pool available during the monsoon months which are them rapidly degraded by microbial communities [44] adapted to these conditions. In contrast, the increase of carbohydrates by the end of the monsoon season (September and August) can result from the increase in surface vegetation that occurs after the monsoon.

Similar to previous studies [45], increased precipitation resulted in an initial increase of labile metabolites, as indicated by the increase in metabolite diversity (Fig. 2f) and NOSC (Supplementary Fig. 2e) at the beginning of the monsoon season. The increase in labile compounds may be derived from the increased vegetation and microbial production and mortality [31] caused by the increased water availability, which fuels microbial metabolism as evidenced by the increase in metabolite transformations among the N- and S-containing metabolites (Supplementary Fig. 2f). The subsequent decrease in metabolite diversity and NOSC by season's end demonstrates efficient microbial processing of labile compounds, highlighting how community-level metabolic responses shape organic matter composition in arid soils.

These metabolomic insights reinforce our dual-level resilience framework by demonstrating how individual metabolic responses collectively shape community resource pools and ecosystem function. The distinct metabolite profiles and their temporal dynamics reflect both community-level adaptation strategies and individual metabolic flexibility in response to environmental fluctuations.

**Supplementary Note 2**

**Carbon and Nitrogen Metabolism Pathways Supporting Microbial Resilience in Arid Soils**

*This note examines the distribution and expression of key metabolic pathways across arid soil microbiomes, focusing on carbon fixation strategies and nitrogen cycling mechanisms. The analysis reveals unexpected complexity in carbon fixation pathways and specialized nitrogen transformation capabilities, particularly in Thermoproteota, demonstrating how metabolic versatility at both individual and community levels enables ecosystem function under extreme conditions. These findings highlight how diverse metabolic strategies contribute to microbial resilience in arid environments, with implications for understanding ecosystem responses to climate change.*

Comprehensive analysis of metabolic potential across recovered genomes revealed sophisticated adaptations for energy generation in arid conditions. Using Kofam [46], SCyc [47], and NCyc [48] databases combined with curated KEGG module definitions [49], we identified functional pathways when at least 50% of required genes were present (Supplementary Fig. 4). This approach provides unprecedented insights into the metabolic capabilities of arid soil microbiomes, revealing novel adaptations to extreme environments.

***Carbon Fixation Pathways***

The Calvin-Benson-Banshan (CBB) cycle, considered the most widespread CO2 uptake pathway in terrestrial ecosystems [50], appeared in twelve bacterial genomes across Actinomycetota, Desulfobacterota, and Pseudomonadota. These MAGs encoded ribulose-1,5-biphosphate carboxylase (RuBisCO) for transforming ribulose-1,5-phosphate to 3-phosphoglycerate [51] (Supplementary Fig. 4). The broad distribution across taxa aligns with previous observations [52]. Notably, some Actinomycetota displayed potential novel energy generation strategies, possibly coupling H2 oxidation through high-affinity hydrogenases with the CBB cycle, suggesting adaptation to resource-limited conditions (Jordaan, 2020). This finding opens new avenues for understanding microbial survival strategies in extreme conditions and may have implications for biogeochemical cycling in arid regions.

Despite previous suggestions favoring energy-conserving pathways like Wood-Ljungdahl, reverse TCA (rTCA), and dicarboxylate/4-hydroxybutyrate (DC/4HB) cycles in oligotrophic environments [50,53]) our analysis revealed more complex patterns. While many MAGs from Acidobacteriota, Actinomycetota, Bacillota, Chloroflexota, Gemmatinomonadota, and Pseudomonadota showed rTCA cycle components for carbon fixation steps - particularly succinyl-CoA to 2-oxoglutarate and 2-oxoglutarate to isocitrate conversion [52], - none encoded complete acetyl-CoA and $CO_2$ to pyruvate conversion pathways. Only one Nitrospirota genome possessed ATP-dependent citrate lyases for cycle completion. Meanwhile, 4 genomes from Thermoproteota were found to encode the gene for the dehydration of 4-hydroxybutyryl-CoA to crotonyl-CoA (Supplementary Fig. 4, a key step in carbon fixation using the 3-Hydroxypropionate/4-Hydroxybutyrate (3HP/4HB) and DC/4HB cycles [54]. Even though none of the Thermoproteota MAGs encode a complete 3HP/4HB or DC/4HB, even the presence of a rudimentary 3HP/4HB cycle may provide this archaea with the ability of coassimilate other organic metabolites [55]. This finding highlights the potential for novel or modified carbon fixation pathways in arid soil microbiomes, warranting further investigation.

***Nitrogen Metabolism***

Our study also sheds light on the nitrogen metabolism of arid soil microbiomes, a critical aspect given the limiting nature of nitrogen in these ecosystems [56]. Notably, we identified nitrifying organisms exclusively within the phylum Thermoproteota, all belonging to the ammonia-oxidizing archaea (AOA) order Nitrososphaerales (Supplementary Fig. 4), an order of ammonia oxidizing archaea (AOA) widely distributed in terrestrial and marine ecosystems [57]. The presence of amoABC genes in these archaea for ammonia oxidation [58] (Fig. 6), is coupled with the absence of hydroxylamine dehydrogenase (HAO), suggesting potential novel routes for hydroxylamine oxidation, possibly involving blue copper proteins or cupredoxin-like proteins [57,59]. This finding not only expands our understanding of nitrification in arid soils but also points to potential new pathways in archaeal nitrogen metabolism.

Furthermore, our analysis revealed the widespread presence of nearly complete pathways for dissimilatory nitrate reduction to ammonia (DNRA) across various phyla, including Acidobacteria, Actynomycetota, Chloroflexota, and Pseudomonadota (Supplementary Fig. 4). While DNRA is known to be ubiquitous in terrestrial environments [60], a process known to depend on C/N ratios, and is more favorable in environments with high C and low nitrogen concentrations [61] its significance in arid systems has been underexplored [56]. Our findings suggest that DNRA may play a crucial role in nitrogen retention in arid soils, particularly under conditions of high carbon and low nitrogen availability. These insights into carbon fixation and nitrogen metabolism in arid soil microbiomes have broad implications. They not only enhance our understanding of microbial adaptations to extreme environments but also provide a foundation for predicting ecosystem responses to climate change in arid regions.

These insights into carbon fixation and nitrogen metabolism demonstrate how specialized metabolic adaptations enable both individual survival and community-level function in extreme conditions, supporting our dual-level resilience framework.

### Supplementary Note 3

#### Dormancy and Stress Protection Mechanisms Across Arid Soil Taxa

*This note examines key survival mechanisms employed by microorganisms in arid soils, focusing on sporulation pathways, carbon storage through glycogen metabolism, and protective polysaccharide production (EPS/LPS). The analysis reveals differential expression patterns of these mechanisms across taxa and temporal conditions, with some groups showing specialized adaptations during wet versus dry periods, highlighting diverse strategies for persistence in extreme conditions.*

Essential actinobacterial sporulation genes, including the cell division regulator WhiA (K09762) and transcriptional regulator WhiB (K18955, K18956, K18957) [62], were encoded across Actinomycetota, Bacillota, Chloroflexota, and Eremiobacteriota genomes. However, expression was detected only in Actinomycetota and Chloroflexota. Similarly, while Bacillota genomes encoded sporulation genes (spoIIB, sda, spoVID, safA), no expression was observed (Fig. 4c, Supplementary Figure 5, Supplementary Table 3).

Genes encoding for the synthesis of glycogen, necessary for entering and reviving from resting stages [63], were found predominantly expressed in wet months in Acidobacteria, Actinomycetota and Pseudomonadote genomes (n = 8). Similarly, genes encoding for glycogen degradation were found expressed for most MAGs (n = 35) in wet moths with the exception of Tectomicrobia, which showed the highest expression compared with the other MAGs in dry months.

In the case of EPS production, only a few genomes of Pseudomonadota were found to encode genes for the biosynthesis of Poly-N-acetyl-glucosamine (PNAG), Pel polysaccharide and Vibrio-like polysaccharides, but several genomes encoded genes for the assembly and transport of LPS. Pel polysaccharide was expressed only in 1 Pseudomonadota genome while LPS was expressed in 7 phyla including, Pseudomonadota , Desulfobacterota, Gemmatimonadota, Methylomirabilota and Nitrospirota.

Bacteria secrete polysaccharides during growth to form protective capsules or regulate their environment, aiding in chemical reactions, trapping nutrients, and mitigating drought or salinity [6465]. While various taxa can produce EPS, studies have identified Pseudomonadota (Proteobacteria) as primary producers of EPS and lipopolysaccharides (LPS) in biocrusts and rhizosphere soils [66,67].

**Supplementary Note 4**

**Thermoproteota Metabolic Capacities in Arid Soils**

*The following detailed metabolic analysis of Thermoproteota demonstrates how individual-level adaptations support community resilience in arid ecosystems through regulated gene expression and metabolic flexibility.*

Nitrogen metabolism

Thermoproteota genomes, belonging to Nitrososphaera, TH5893 and of unknown genera, possessed orthologous genes for denitrification pathway, including NirK (K00368), norB (K04561) and nosZ (K00376), but none of them encode for narG (K00370) or napA (K02567) for the reduction of nitrate into nitrite (Supplementary Table 5). All these were found to be expressed in the metatranscriptomics dataset, with the higher expression of the NirK gene observed in three Nitrososphaera genomes during May and July. It has been suggested that NirK is essential for ammonia oxidation in ammonia oxidizing archaea (AOA) as NO (product of NirK) acts as an intermediate [68] , however its precise function remains unclear [58]. norB and nosZ genes have been described in other Thermoproteota genera such as Nitrosocosmicus but not in Nitrososphaera or TH5893 [58].

Five genomes encoded genes NR (K10534) for nitrate reduction via the assimilatory nitrate reduction pathway, but three of them also encoding the nirA gene (K00366) for the reduction of nitrite to ammonia. Both those genes were found to be expressed with the nirA gene having a higher expression during the dry months. Within the dissimilatory nitrate reduction pathway, most Thermoproteota genomes (n = 7) encoded only for the nirD gene (K00363) for the transformation of nitrite to ammonia in a complex with nirB (K00362), which was found only in two genomes. The highest expression of nirD was observed during dry months in three Nitrososphaera genomes.

The expression of glutamate dehydrogenase, gudB, rocG (K00260) and glutamate dehydrogenase NAD(P)+, GLUD1_2 (K00261) involved in the interconversion of glutamate to 2-oxoglutarate and ammonia was expressed as well, the former expressed only in one Nitrososphaera genome with highest expression observed in dry months. While the expression of glutamine synthetase (K01915) that couples ammonium assimilation to glutamate synthesis was observed at lower levels and expressed in other genomes. These suggest that Nitrososphaera genomes can obtain nitrogen from different organic molecules as observed in other N-limiting environments such as geothermal springs [69].

Sulfur metabolism

Genes involved in the assimilatory sulfate reduction pathway were found to be encoded and expressed including sat (K00958), cysC (K00860), cysH (K00390), cysI (K00381) and sir (K00392). Also, genes encoding the sulfonate transport system, the substrate-binding protein ssuA (K15553) and the ATP-binding protein ssuB (K15555) were expressed as well. The expression of sulfide:quinone oxidoreductase (K17218) for the oxidation of sulfide to zero-valent sulfur was observed as well as thiosulfate sulfurtransferase glpE (K02439) involved in the transformation of thiosulfate to sulfite. This indicates that Nitrososphaera genomes are able to obtain sulfur from organic and inorganic compounds, no major changes in the expression of these genes were observed through the monsoon, although the expression levels were low.

Carbon fixation

Incomplete carbon fixation pathways were found encoded in the Thermoproteota genomes. For the Dicarboxylate-hydroxybutyrate (DC/4HB) pathway we found the presence and expression of 4-hydroxybutyryl-CoA dehydratase / vinylacetyl-CoA-Delta-isomerase (K14534), a key enzyme of this pathway. Other genes encoded and expressed were enoyl-CoA hydratase / 3-hydroxyacyl-CoA dehydrogenase (K15016), succinyl-CoA synthetase alpha subunit (K01902) and the succinate dehydrogenase flavoprotein subunit (K00239) and the succinate dehydrogenase iron-sulfur subunit (K00240). However other key enzymes of this pathway, succinic semialdehyde reductase (NADPH) (K14465) and 4-hydroxybutyrate-CoA ligase (K14467,K18861,K25774) were not encoded by any genome. This could be due to the lack of metagenome sequencing coverage, complete DC/4HB has been described in other Thermorproteota families different than Nitrososphaeracea [69].

On the other hand, the Hydroxypropionate-hydroxybutylate (3-HP-HB) pathway was almost completely encoded and expressed. The abfD (K14534) gene involved in both pathways, DC/4HB and 3HP/4HB, showed the highest expression during dry months (May and October) in 3 Nisotrosphaera genome and one genome of unknown genus.

Within the Reductive citrate cycle, the following genes were found to be encoded and expressed; pyruvate, orthophosphate dikinase (K01006), succinyl-CoA synthetase alpha subunit (k01902), the succinate dehydrogenase flavoprotein subunit (K00239) and the succinate dehydrogenase iron-sulfur subunit (K00240), the 2-oxoglutarate/2-oxoacid ferredoxin oxidoreductase subunits alpha (K00174) and beta (K00175). The highest expression was observed in dry months for the later genes encoding the transformation of succinyl-CoA to 2-oxoglutarate.

Carbon metabolism

Central carbon metabolism

Most of the reactions within the Glycolysis/gluneogenesis pathway were found encoded and expressed. The highest expression was observed in dry months for the fructose 1,6-bisphosphate aldolase/phosphatase (K01622) gene encoding the transformation of fructose-6P to glyceraldehyde-3P and pyruvate kinase (K00873) encoding the transformation of phosphoenolpyruvate to pyruvate.

Within the Citrate cycle, only the genes involved in the 3 first reactions of the second carbon oxidation pathway were encoded and expressed.

Almost all of the non-oxidative phase of the pentose phosphate pathway was encoded and expressed with the exception of the transformation of xylulose-5P to ribulose-5P. In general, all the steps within this pathway seemed to be more expressed during dry months, the higher expression was observed in the ribose 5-phosphate isomerase A gene (K01807) involved in the transformation of ribulose-5P to ribose-5P. The gene (K00033), part of the oxidative phase, encoding the transformation of 6-phospho-D-gluconate to ribulose-5P was also expressed in 3 Nitrososphaera genomes.

Also the ribose-phosphate pyrophosphokinase genes (K00948) that encodes the enzyme catalyzing the biosynthesis of 5-Phospho-alpha-D-ribose 1-diphosphate (PRPP) was expressed in 7 Thermoproteota genomes, 3 Nitrososphaera genomes showed the highest expression in dry months.

Other metabolism

Genes involved in three reactions within the Formaldehyde assimilation, serine pathway were found to be encoded and expressed including glycine hydroxymethyltransferase(K00600), glycerate 2-kinase (K11529) and enolase 1/2/3 (K10689).

Amino acid metabolism

Three Thermoproteota MAGs encoded the enzymes for the glycine cleavage system (and other 3 genomes have an incomplete pathway), but only a single Nitrososphaera genome expressed all reactions within this pathway, while other two genomes expressed only the first and last reaction. The genes encoding the glycine cleavage system P protein (glycine dehydrogenase) subunits 1 (K00282) and 2 (K00283), that are involved in the transformation of glycine to $CO_2$, were more expressed during dry months in 3 Nitrososphaera genomes. Meanwhile, the glycine cleavage system T protein (aminomethyltransferase)(K00605), involved in the transformation of

Tetrahydrofolate to Ammonia was only expressed in one genomes with a higher expression observed in July. The dihydrolipoyl dehydrogenase gene(K00382), involved in the transformation of diihydrolipoylprotein to lipoylprotein, was more expressed during dry months in 4 genomes.

Genes for the two reactions for the transformation of 3-phospho-D-glycerate to serine within the biosynthesis of the serine pathway, encoded by the D-3-phosphoglycerate dehydrogenase / 2-oxoglutarate reductase (K00058) and phosphoserine phosphatase (K01079) respectively, were found encoded in most Thermoproteota genomes (n=6), but they were only also expressed in four of them.

Cofactors

These archaea also encoded genes within the sulfur relay pathway including moaD (K03636), moeB (K21029), iscS (K04487), moaE (K03635) and moaA (K03639) for the biosynthesis of molybdenum cofactor (MOCO). This has been previously reported and suggested to provide additional protection for oxidative stress [58]

All genes involved in the redox cofactor, coenzyme F420 were expressed as well. It has been suggested that this widely distributed cofactor confers advantages to soil microbial communities by allowing them to detoxify, decompose and synthesize a broad range of organic compounds [70].

Genes involved in the aerobic and anaerobic cobalamin (vitamin $B_{12}$) biosynthesis were expressed as well. Genomic analysis indicated that Thermoproteota is an important cobalamin producer in marine and soil environments [71,72].

### REFERENCES


1. Woodcroft, B. J. *et al.* SingleM and Sandpiper: Robust microbial taxonomic profiles from metagenomic data. *bioRxiv* 2024.01.30.578060 (2024) doi:10.1101/2024.01.30.578060.
2. Kushwaha, P. *et al.* Arid Ecosystem Vegetation Canopy-Gap Dichotomy: Influence on Soil Microbial Composition and Nutrient Cycling Functional Potential. *Appl. Environ. Microbiol.* 87, (2021).
3. McHugh, T. A. *et al.* Climate controls prokaryotic community composition in desert soils of the southwestern United States. *FEMS Microbiol. Ecol.* 93, fix116 (2017).
4. Idris, H., Goodfellow, M., Sanderson, R., Asenjo, J. A. & Bull, A. T. Actinobacterial rare biospheres and dark matter revealed in habitats of the Chilean Atacama Desert. *Sci. Rep.* 7, 8373 (2017).
5. Belov, A. A., Cheptsov, V. S. & Vorobyova, E. A. Soil bacterial communities of Sahara and Gibson deserts: Physiological and taxonomical characteristics. *AIMS Microbiol.* 4, 685–710 (2018).
6. Mawar, R., Ranawat, M., Sharma, S. K. & Sayyed, R. Z. Exploring microbial diversity of arid regions of globe for agricultural sustainability: A revisit. in *Plant Growth Promoting Microorganisms of Arid Region* 1–25 (Springer Nature Singapore, Singapore, 2023).
7. Demergasso, C. *et al.* Hyperarid soil microbial community response to simulated rainfall. *Front. Microbiol.* 14, 1202266 (2023).
8. Zhang, Z. *et al.* Prokaryotic taxonomy and functional diversity assessment of different sequencing platform in a hyper-arid Gobi soil in Xinjiang Turpan Basin, China. *Front. Microbiol.* 14, 1211915 (2023).
9. Neilson, J. W. *et al.* Life at the hyperarid margin: novel bacterial diversity in arid soils of the Atacama Desert, Chile. *Extremophiles* 16, 553–566 (2012).
10. Chanal, A. *et al.* The desert of Tataouine: an extreme environment that hosts a wide diversity of microorganisms and radiotolerant bacteria. *Environ. Microbiol.* 8, 514–525 (2006).
11. Hu, D., Zang, Y., Mao, Y. & Gao, B. Identification of molecular markers that are specific to the class Thermoleophilia. *Front. Microbiol.* 10, 1185 (2019).
12. Xie, F. & Pathom-Aree, W. Actinobacteria from desert: Diversity and biotechnological applications. *Front. Microbiol.* 12, 765531 (2021).
13. Gao, Q. & Garcia-Pichel, F. Microbial ultraviolet sunscreens. *Nat. Rev. Microbiol.* 9, 791–802 (2011).
14. Davis, K. E. R., Sangwan, P. & Janssen, P. H. Acidobacteria, Rubrobacteridae and Chloroflexi are abundant among very slow-growing and mini-colony-forming soil bacteria: Slow-growing and mini-colony-forming soil bacteria. *Environ. Microbiol.* 13, 798–805 (2011).
15. Schimel, J., Balser, T. C. & Wallenstein, M. Microbial stress-response physiology and its implications for ecosystem function. *Ecology* 88, 1386–1394 (2007).
16. Fierer, N. *et al.* Cross-biome metagenomic analyses of soil microbial communities and their functional attributes. *Proc. Natl. Acad. Sci. U. S. A.* 109, 21390–21395 (2012).
17. Hwang, Y. *et al.* Leave no stone unturned: individually adapted xerotolerant Thaumarchaeota sheltered below the boulders of the Atacama Desert hyperarid core. *Microbiome* 9, 234 (2021).
18. Lian, W.-H. *et al.* Culturomics- and metagenomics-based insights into the microbial community and function of rhizosphere soils in Sinai desert farming systems. *Environ. Microbiome* 18, 4 (2023).
19. Liu, Q., Chen, Y. & Xu, X.-W. Genomic insight into strategy, interaction and evolution of nitrifiers in metabolizing key labile-dissolved organic nitrogen in different environmental

niches. *Front. Microbiol.* 14, 1273211 (2023).
20. Ray, A. E. *et al.* Atmospheric chemosynthesis is phylogenetically and geographically widespread and contributes significantly to carbon fixation throughout cold deserts. *ISME J.* 16, 2547–2560 (2022).
21. Karner, M. B., DeLong, E. F. & Karl, D. M. Archaeal dominance in the mesopelagic zone of the Pacific Ocean. *Nature* 409, 507–510 (2001).
22. Xu, L. *et al.* Genome-resolved metagenomics reveals role of iron metabolism in drought-induced rhizosphere microbiome dynamics. *Nat. Commun.* 12, 3209 (2021).
23. Tolić, N. *et al.* Formularity: Software for automated formula assignment of natural and other organic matter from ultrahigh-resolution mass spectra. *Anal. Chem.* 89, 12659–12665 (2017).
24. Wilson, R. M. *et al.* Soil metabolome response to whole-ecosystem warming at the Spruce and Peatland Responses under Changing Environments experiment. *Proc. Natl. Acad. Sci. U. S. A.* 118, (2021).
25. Wilson, R. M. *et al.* Plant organic matter inputs exert a strong control on soil organic matter decomposition in a thawing permafrost peatland. *Sci. Total Environ.* 820, 152757 (2022).
26. AminiTabrizi, R. *et al.* Controls on Soil Organic Matter Degradation and Subsequent Greenhouse Gas Emissions Across a Permafrost Thaw Gradient in Northern Sweden. *Front Earth Sci. Chin.* 8, (2020).
27. Hildebrand, G. A. *et al.* Uncovering the dominant role of root metabolism in shaping rhizosphere metabolome under drought in tropical rainforest plants. *Sci. Total Environ.* 899, 165689 (2023).
28. Lin, Y. *et al.* Differential effects of redox conditions on the decomposition of litter and soil organic matter. *Biogeochemistry* 154, 1–15 (2021).
29. Sheng, M. *et al.* Spatial and molecular variations in forest topsoil dissolved organic matter as revealed by FT-ICR mass spectrometry. *Sci. Total Environ.* 895, 165099 (2023).
30. Qi, Y. *et al.* Assessment of molecular diversity of lignin products by various ionization techniques and high-resolution mass spectrometry. *Sci. Total Environ.* 713, 136573 (2020).
31. Tfaily, M. M., Hess, N. J., Koyama, A. & Evans, R. D. Elevated [CO2] changes soil organic matter composition and substrate diversity in an arid ecosystem. *Geoderma* 330, 1–8 (2018).
32. Ramond, J.-B. & Cowan, D. A. Microbial ecology of hot desert soils. in *Ecological Studies* 89–110 (Springer International Publishing, Cham, 2022).
33. Yu, Z. *et al.* Molecular insights into the transformation of dissolved organic matter during hyperthermophilic composting using ESI FT-ICR MS. *Bioresour. Technol.* 292, 122007 (2019).
34. Valle, J. *et al.* Extensive processing of sediment pore water dissolved organic matter during anoxic incubation as observed by high-field mass spectrometry (FTICR-MS). *Water Res.* 129, 252–263 (2018).
35. Bao, H., Michalski, G. M. & Thiemens, M. H. Sulfate oxygen-17 anomalies in desert varnishes. *Geochim. Cosmochim. Acta* 65, 2029–2036 (2001).
36. Moreno-Rodríguez, V. *et al.* Historical trends and sources of TSP in a Sonoran desert city: Can the North America Monsoon enhance dust emissions? *Atmos. Environ. (1994)* 110, 111–121 (2015).
37. Gu, A. & Eastoe, C. J. The origins of sulfate in Cenozoic non-marine evaporites in the Basin and-range province, southwestern North America. *Geosciences (Basel)* 11, 455 (2021).
38. Su, S. *et al.* High molecular diversity of organic nitrogen in urban snow in North China. *Environ. Sci. Technol.* 55, 4344–4356 (2021).
39. Gonsior, M. *et al.* Molecular characterization of effluent organic matter identified by ultrahigh resolution mass spectrometry. *Water Res.* 45, 2943–2953 (2011).

40. Tfaily, M. M. *et al.* Vertical stratification of peat pore water dissolved organic matter composition in a peat bog in northern Minnesota: Pore water DOM composition in a peat bog. *J. Geophys. Res. Biogeosci.* 123, 479–494 (2018).
41. Ma, T. *et al.* Divergent accumulation of microbial necromass and plant lignin components in grassland soils. *Nat. Commun.* 9, 3480 (2018).
42. Šťovíček, A., Kim, M., Or, D. & Gillor, O. Microbial community response to hydration-desiccation cycles in desert soil. *Sci. Rep.* 7, 45735 (2017).
43. Brangarí, A. C., Manzoni, S. & Rousk, J. The mechanisms underpinning microbial resilience to drying and rewetting – A model analysis. *Soil Biol. Biochem.* 162, 108400 (2021).
44. Chen, H. *et al.* Molecular insights into Arctic soil organic matter degradation under warming. *Environ. Sci. Technol.* 52, 4555–4564 (2018).
45. Chen, Q., Niu, B., Hu, Y., Luo, T. & Zhang, G. Warming and increased precipitation indirectly affect the composition and turnover of labile-fraction soil organic matter by directly affecting vegetation and microorganisms. *Sci. Total Environ.* 714, 136787 (2020).
46. Aramaki, T. *et al.* KofamKOALA: KEGG Ortholog assignment based on profile HMM and adaptive score threshold. *Bioinformatics* 36, 2251–2252 (2020).
47. Yu, X. *et al.* SCycDB: A curated functional gene database for metagenomic profiling of sulphur cycling pathways. *Mol. Ecol. Resour.* 21, 924–940 (2021).
48. Tu, Q., Lin, L., Cheng, L., Deng, Y. & He, Z. NCycDB: a curated integrative database for fast and accurate metagenomic profiling of nitrogen cycling genes. *Bioinformatics* 35, 1040–1048 (2019).
49. Kanehisa, M. The KEGG database. *Novartis Found. Symp.* 247, 91–101; discussion 101–3, 119–28, 244–52 (2002).
50. Yang, Y. *et al.* Changes in soil microbial carbon fixation pathways along the oasification process in arid desert region: A confirmation based on metagenome analysis. *Catena* 239, 107955 (2024).
51. Tabita, F. R. *et al.* Function, structure, and evolution of the RubisCO-like proteins and their RubisCO homologs. *Microbiol. Mol. Biol. Rev.* 71, 576–599 (2007).
52. Garritano, A. N., Song, W. & Thomas, T. Carbon fixation pathways across the bacterial and archaeal tree of life. *PNAS Nexus* 1, gac226 (2022).
53. Jiang, Q., Jing, H., Jiang, Q. & Zhang, Y. Insights into carbon-fixation pathways through metagonomics in the sediments of deep-sea cold seeps. *Mar. Pollut. Bull.* 176, 113458 (2022).
54. Berg, I. A. Ecological aspects of the distribution of different autotrophic CO2 fixation pathways. *Appl. Environ. Microbiol.* 77, 1925–1936 (2011).
55. Hatzenpichler, R. Diversity, physiology, and niche differentiation of ammonia-oxidizing archaea. *Appl. Environ. Microbiol.* 78, 7501–7510 (2012).
56. Ramond, J.-B., Jordaan, K., Díez, B., Heinzelmann, S. M. & Cowan, D. A. Microbial Biogeochemical Cycling of Nitrogen in Arid Ecosystems. *Microbiol. Mol. Biol. Rev.* 86, e00109–21 (2022).
57. Stahl, D. A. & de la Torre, J. R. Physiology and diversity of ammonia-oxidizing archaea. *Annu. Rev. Microbiol.* 66, 83–101 (2012).
58. Bei, Q. *et al.* Metabolic potential of Nitrososphaera-associated clades. *ISME J.* 18, (2024).
59. Diamond, S. *et al.* Soils and sediments host Thermoplasmata archaea encoding novel copper membrane monooxygenases (CuMMOs). *ISME J.* 16, 1348–1362 (2022).
60. Yang, W. H., Ryals, R. A., Cusack, D. F. & Silver, W. L. Cross-biome assessment of gross soil nitrogen cycling in California ecosystems. *Soil Biol. Biochem.* 107, 144–155 (2017).
61. Pandey, C. B. *et al.* DNRA: A short-circuit in biological N-cycling to conserve nitrogen in terrestrial ecosystems. *Sci. Total Environ.* 738, 139710 (2020).
62. Guiza Beltran, D., Wan, T. & Zhang, L. WhiB-like proteins: Diversity of structure, function

and mechanism. *Biochim. Biophys. Acta Mol. Cell Res.* 1871, 119787 (2024).
63. Nariya, H. & Inouye, S. An effective sporulation of Myxococcus xanthus requires glycogen consumption via Pkn4-activated 6-phosphofructokinase: Regulation of glycogen metabolism by PSTK in M. xanthus. *Mol. Microbiol.* 49, 517–528 (2003).
64. Meier, D. V., Imminger, S., Gillor, O. & Woebken, D. Distribution of mixotrophy and desiccation survival mechanisms across microbial genomes in an arid biological soil crust community. *mSystems* 6, (2021).
65. Costa, O. Y. A., Raaijmakers, J. M. & Kuramae, E. E. Microbial extracellular polymeric substances: Ecological function and impact on soil aggregation. *Front. Microbiol.* 9, 1636 (2018).
66. Cania, B. *et al.* Biological soil crusts from different soil substrates harbor distinct bacterial groups with the potential to produce exopolysaccharides and lipopolysaccharides. *Microb. Ecol.* 79, 326–341 (2020).
67. Marasco, R. *et al.* Rhizosheath-root system changes exopolysaccharide content but stabilizes bacterial community across contrasting seasons in a desert environment. *Environ. Microbiome* 17, 14 (2022).
68. Kozlowski, J. A., Stieglmeier, M., Schleper, C., Klotz, M. G. & Stein, L. Y. Pathways and key intermediates required for obligate aerobic ammonia-dependent chemolithotrophy in bacteria and Thaumarchaeota. *ISME J.* 10, 1836–1845 (2016).
69. Qi, Y.-L. *et al.* Analysis of nearly 3000 archaeal genomes from terrestrial geothermal springs sheds light on interconnected biogeochemical processes. *Nat. Commun.* 15, 4066 (2024).
70. Ney, B. *et al.* The methanogenic redox cofactor F420 is widely synthesized by aerobic soil bacteria. *ISME J.* 11, 125–137 (2017).
71. Doxey, A. C., Kurtz, D. A., Lynch, M. D. J., Sauder, L. A. & Neufeld, J. D. Aquatic metagenomes implicate Thaumarchaeota in global cobalamin production. *ISME J.* 9, 461–471 (2015).
72. Lu, X., Heal, K. R., Ingalls, A. E., Doxey, A. C. & Neufeld, J. D. Metagenomic and chemical characterization of soil cobalamin production. *ISME J.* 14, 53–66 (2020).
