## Supplementary Fig for "Stochastic Assembly and Metabolic Network Reorganization Drive Microbial Resilience in Arid Soils"

### SUPPLEMENTARY FIGURES

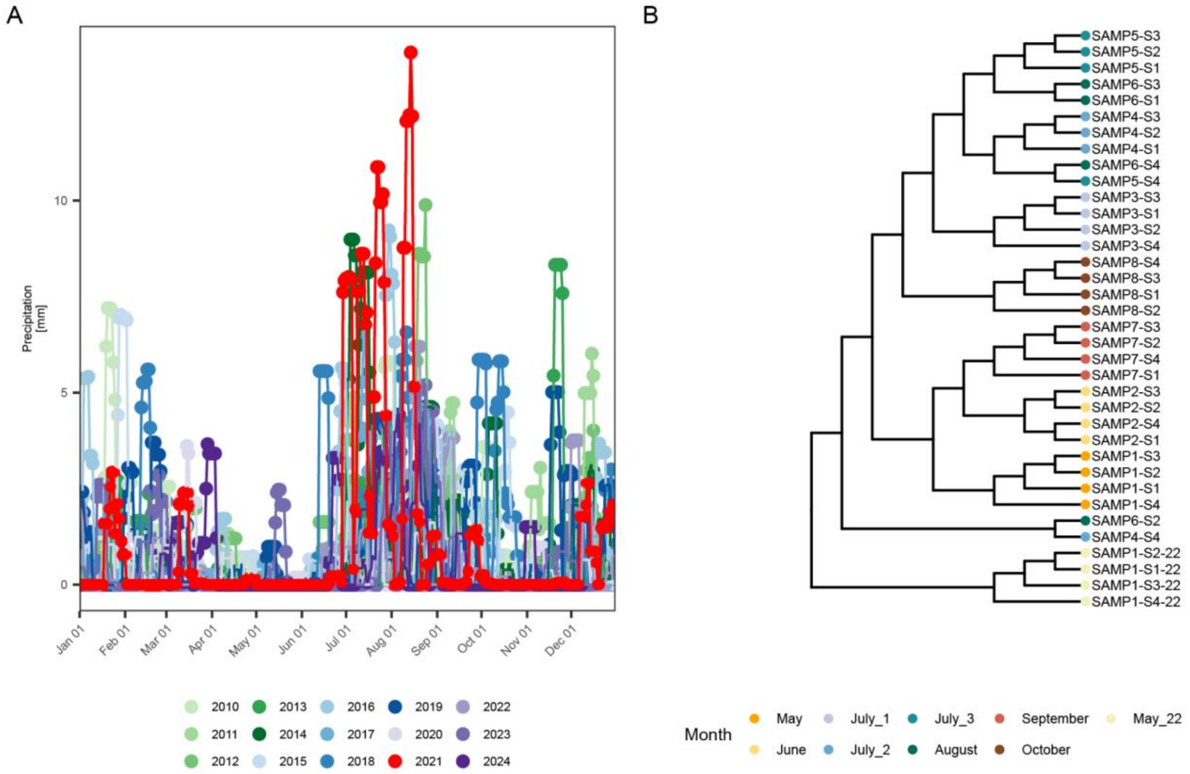


**Supplementary Figure 1.** (A) Seven-day rolling mean of daily precipitation values measured at the AZ_Tucson_11_W station from the period between 2010 - 2024. (B) Hierarchical clustering of all the collected samples based on the measured environmental variables.

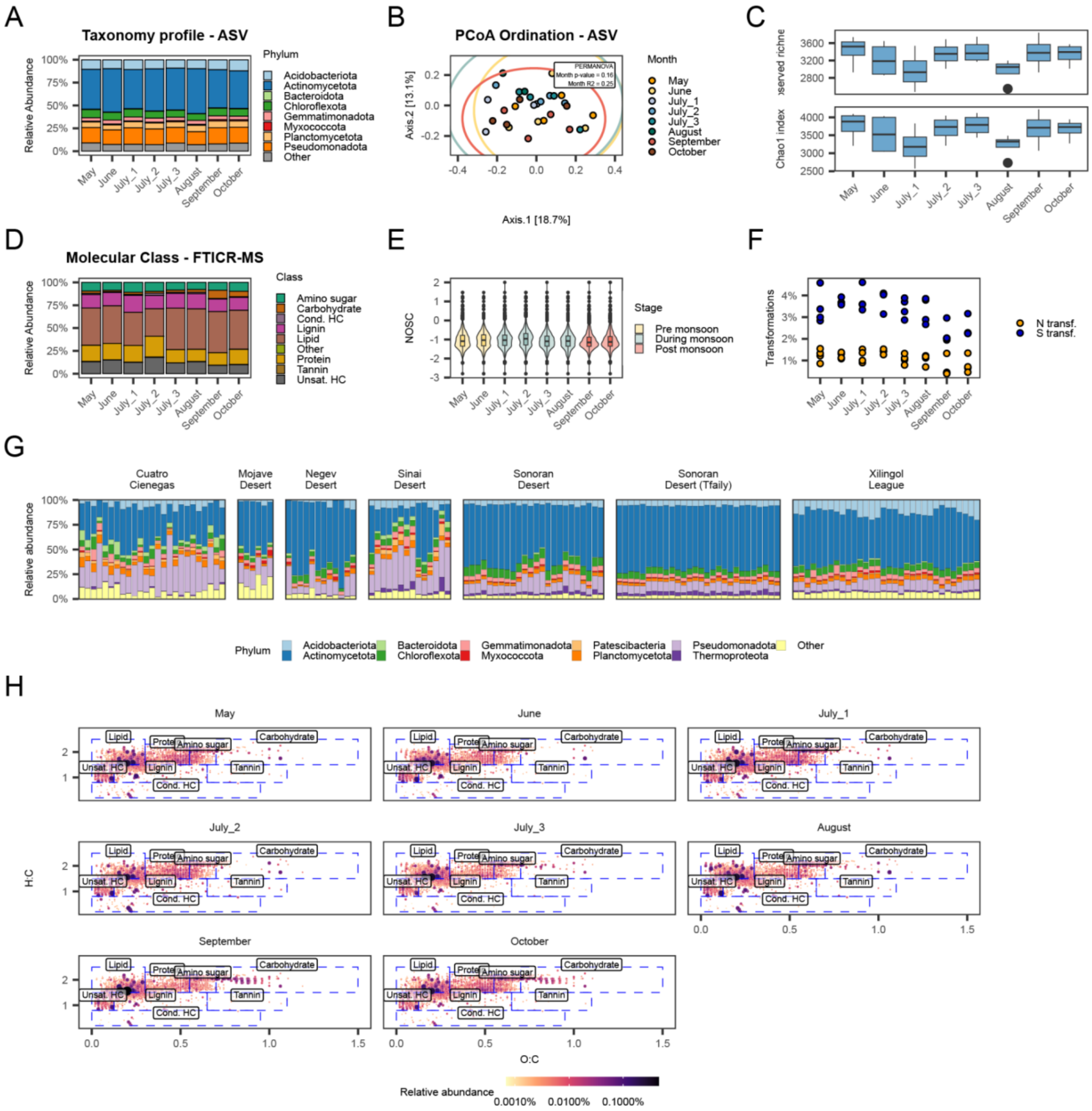


**Supplementary Figure 2.** (A) Average relative abundance of bacterial phyla from ASVs inferred with DADA2 per sampling time (n = 4). (B) Principal component analysis (PCoA) of amplicon sequencing data (n = 32). (C) Alpha diversity indices (observed and Chao1 richness) from ASV relative abundances. (D) Average relative abundance of molecular classes from FTICR-MS positive mode data per sampling time (n = 4). (E) Metabolites' nominal oxidation state of carbon (NOSC) values across sampling points. For box plots (C, E): boxes show upper and lower quartiles with median line; whiskers extend to 1.5 times the interquartile range; points beyond whiskers represent outliers. (F) Number of N- and S-containing compound transformations per sample. (G) Taxonomic profiles of multiple arid ecosystems worldwide. (H) Van Krevelen diagrams showing metabolites at each sampling point, with point color and size indicating relative abundance.

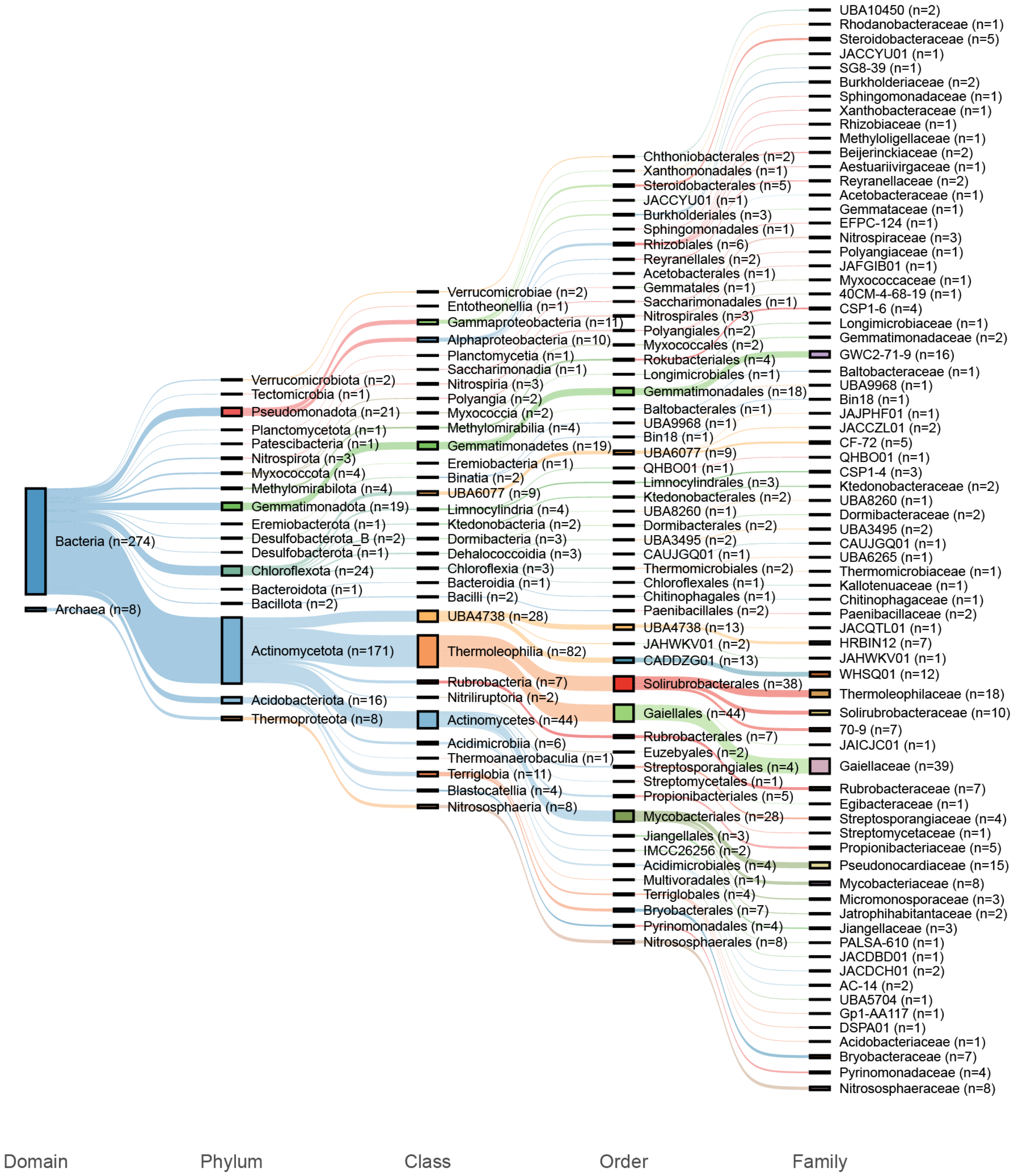


**Supplementary Figure 3.** Sankey diagram showing the taxonomic classification of the 282 medium- and high-quality MAGs generated during this study.

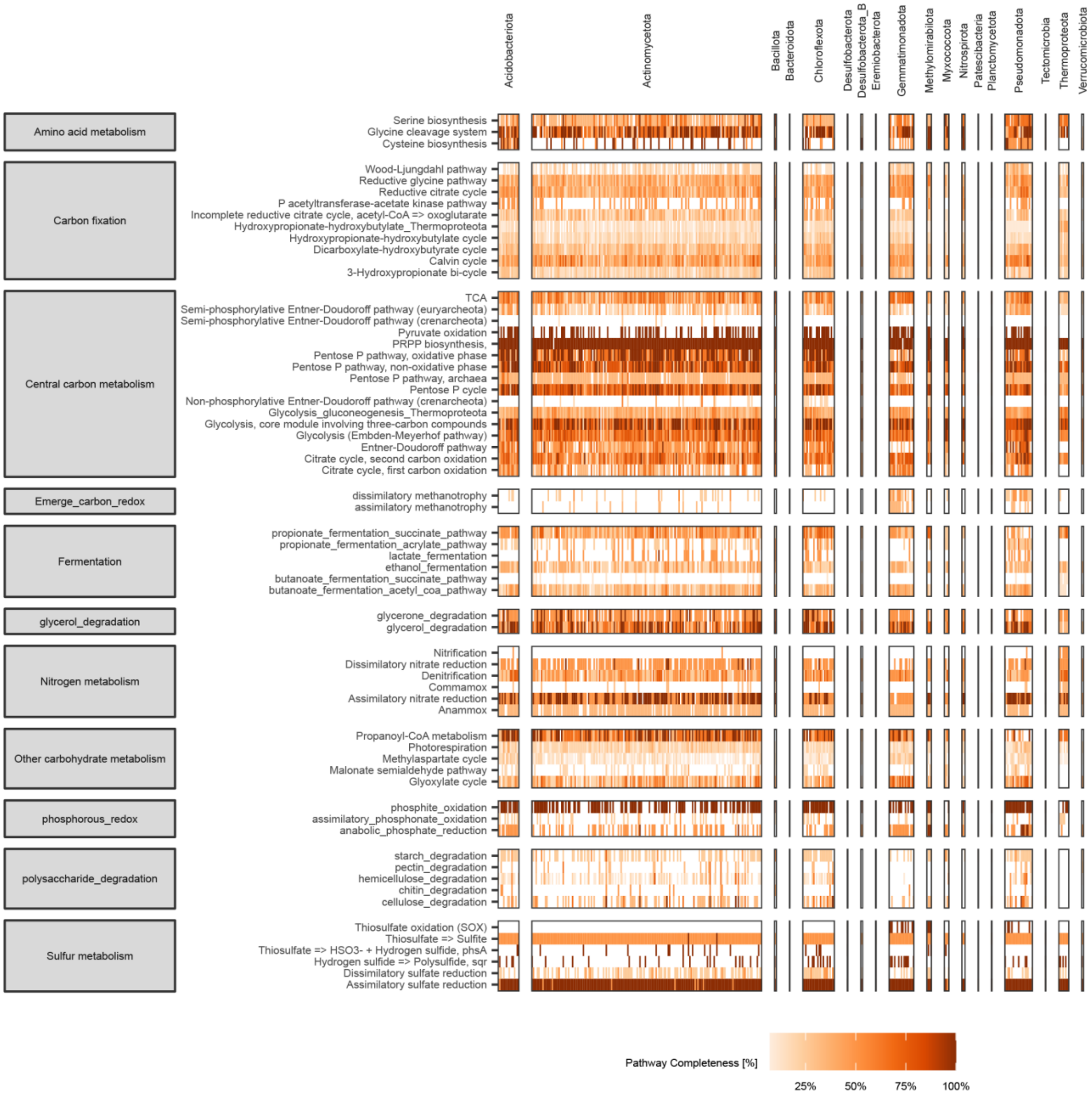


**Supplementary Figure 4.** Heatmap showing the percentage of completeness per MAG of metabolic pathways associated with carbon, nitrogen and sulfur metabolism.

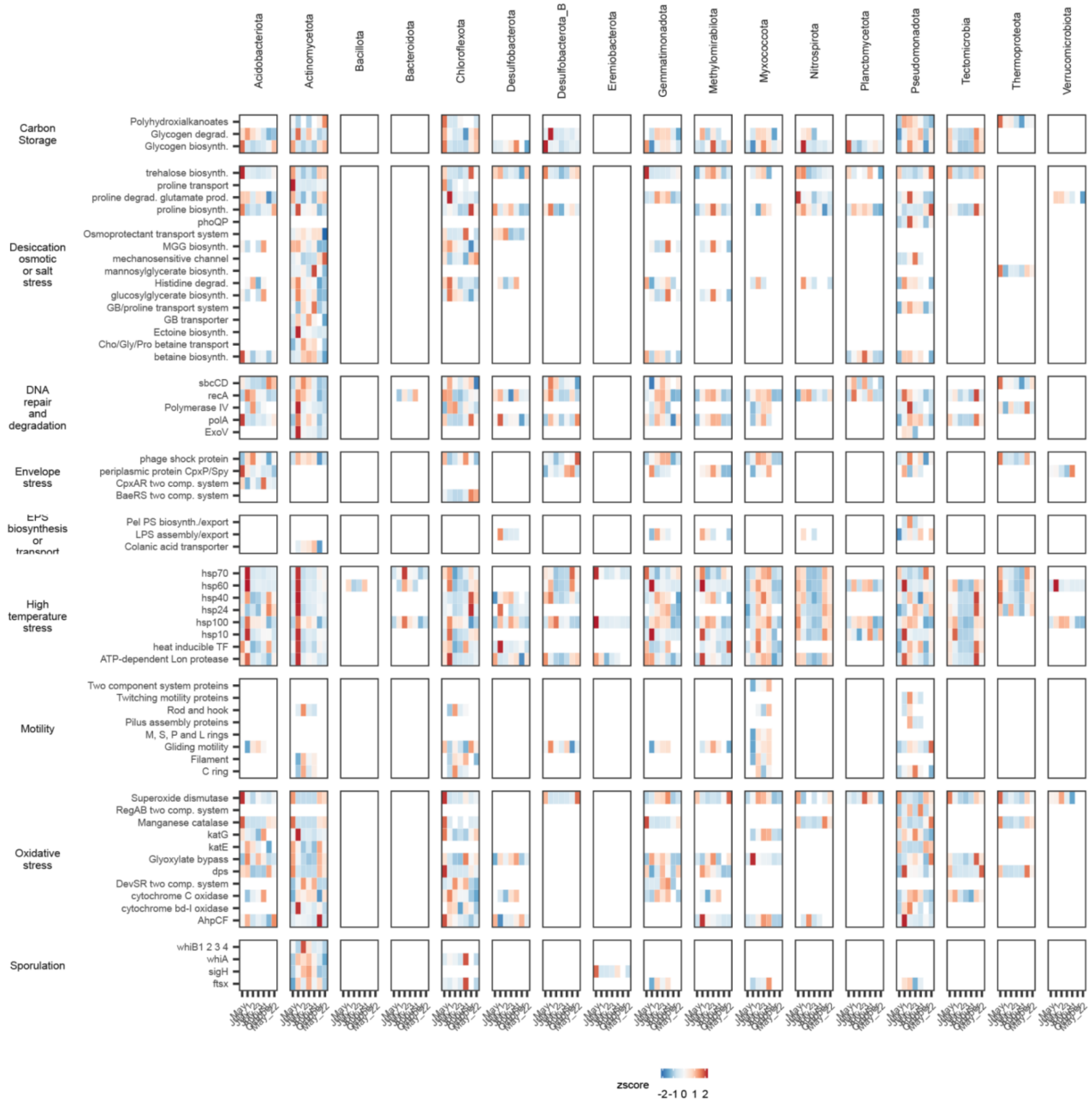


**Supplementary Figure 5.** Heatmap showing the levels of gene expression, agglomerated at phylum level, at different sampling times for various stress tolerance mechanisms defined in Supplementary Table 7. Gene expression was transformed into a z-score for each of the phyla to better represent the changes across the monsoon season.

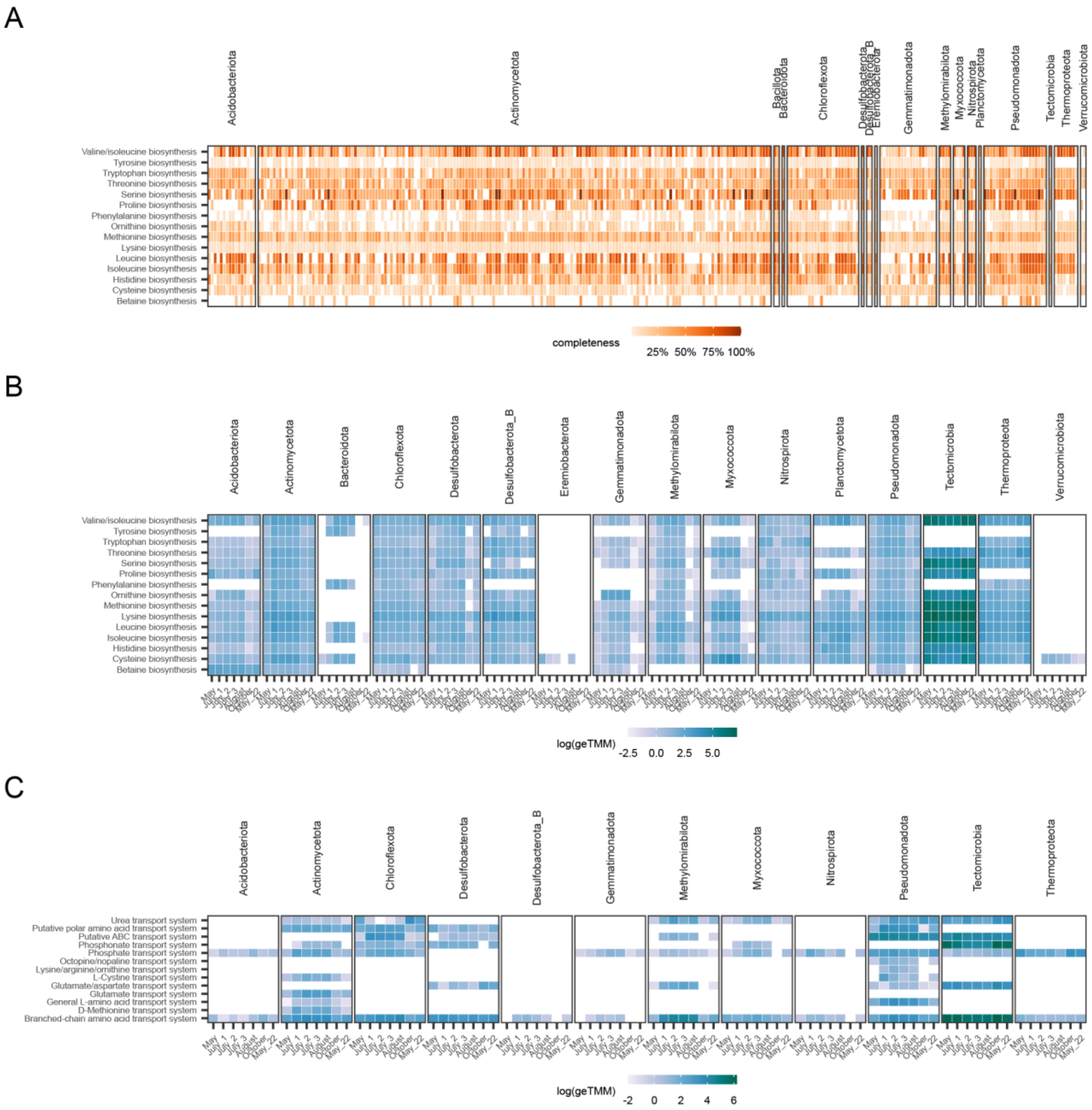


**Supplementary Figure 6.** (A) Heatmap showing the percentage of completeness of different amino acid biosynthetic pathways in each MAG. (B) Heatmap showing the logarithm of gene expression (in geTMM) of different amino acid biosynthetic pathways in each MAG across the different sampling points. (C) Heatmap showing the logarithm of gene expression (in geTMM) of different amino acid transporter systems in each MAG across the different sampling points.
